# An Adeno-Associated Viral vector encoding Neurotrophin 3 injected into affected forelimb muscles modestly improves sensorimotor function after contusive mid-cervical spinal cord injury

**DOI:** 10.1101/2021.02.24.432676

**Authors:** Jared D. Sydney Smith, Vanessa Megaro, Aline Barroso Spejo, Lawrence D. F. Moon

**Author notes:** Corresponding author: Dr Lawrence Moon, Neurorestoration Group, Wolfson Centre for Age-Related Diseases, 16 – 18 Newcomen Street, London, SE1 1UL, United Kingdom. The authors declare that no conflict of interest exists.

## Abstract

Traumatic spinal cord injury (SCI) in humans occurs most frequently in the cervical spine where it can cause substantial sensorimotor impairments to upper limb function. The altered input to spinal circuits below the lesion leads to maladaptive reorganisation which often leads to hyperreflexia in proprioceptive circuits. Neurotrophin 3 (NT3) is growth factor essential for the development of proprioceptive neurons. We have previously shown that following bilateral corticospinal tract axotomy, intramuscular delivery of an Adeno-Associated Viral vector encoding NT3 (AAV-NT3) induces proprioceptive circuit reorganisation linked to functional recovery. To assess its therapeutic effects following a clinically relevant bilateral C5-C6 contusion in rats, AAV-NT3 was injected intramuscularly into the dominant limb 24 hours after injury and forelimb function was assessed over 13 weeks. The injury generated hyperreflexia of a distal forelimb proprioceptive circuit. There was also loss of fine motor skills during reach-and-grasp and walking on a horizontal ladder. Ex vivo magnetic resonance imaging (MRI) revealed atrophy of the spinal cord and white matter disruption throughout the lesion site together with extensive loss of grey matter. Unexpectedly, animals treated with AAV-NT3 had a slightly smaller lesion in the regions close to the epicentre compared to PBS treated animals. Rats treated with AAV-NT3 showed subtly better performance on the horizontal ladder and transient benefits on reach-and-grasp. AAV-NT3 did not normalise hyperreflexia in a treated muscle. The treatment increased the amount of NT3 in treated muscles but, unexpectedly, serum levels were only elevated in a small subset of animals. These results show that this dose and delivery of AAV-NT3 may generate subtle improvements in locomotion but additional treatments will be required to overcome the widespread sensorimotor deficits caused by contusion injury.

## Introduction

### Neuroplasticity drives spontaneous recovery after traumatic spinal cord injury

Traumatic injury to the spinal cord causes axotomy and neuronal death leading to impaired transmission of signals through the lesion site. Damaged neurons have limited capacity to regenerate axons, and as a result the relatively intact spinal circuitry below the lesion is left with reduced or no direct descending control. Despite this it is common for some degree of spontaneous recovery to occur ^1–3^. In response to reduced supraspinal input, the circuitry undergoes a programme of sprouting. This includes increases in afferent input onto motor and premotor neurons ^4–6^ and the formation of relay connections through sprouting of descending fibres that contact long propriospinal neurons running contralateral to the lesion ^7, 8^. Additionally, spared descending axons which remained intact after injury are influenced by the new cellular environment and collateralise to the denervated grey matter ^9, 10^.

This reorganisation of the spared circuitry requires continual local proprioceptive afferent input from muscle spindles, the sense organ in the muscle responsible for detecting stretches within the muscle ^11^. In mice which had proprioceptive neurons ablated, and therefore have abnormal Ia afferent input to the spinal cord, the spontaneous recovery often seen after lateral hemisection does not occur and functional benefits such as regaining plantar stepping and interlimb coordination are absent^6^. These studies highlight the importance of proprioceptive afferent input not only for contributing to spontaneous restoration of function following injury but also their role in maintaining this function after the initial recovery phase^6^.

### Maladaptive plasticity can create abnormally excitable reflexes

Not all reorganisation within the spinal circuitry is beneficial. For those patients with some residual function the development of spasticity, which is thought to occur partly due to maladaptive plasticity, can hamper recovery and impact on post injury health ^12^. Spasticity is an upper motor syndrome characterised traditionally as increased stretch reflexes which are velocity dependent ^13^, which manifest clinically as increased muscle tone and exaggerated reflexes below the neurological level of injury. The syndrome is a chronic condition with symptoms beginning several months after the initial injury and is present in up to 75% of patients suffering symptoms one year after injury ^14, 15^.

Despite being only one component of the spasticity syndrome, hyperreflexia of the stretch reflex is a well-studied phenomenon in both patients and numerous preclinical models of CNS injury ^16–18^. The stretch reflex arc acts to contract the stretching muscle and comprises muscle spindles, Ia afferents, the monosynaptic connection between Ia afferents and homonymous α-motor neuron, and finally the α-motor neuron itself which innervates the muscle at the motor end plate ^19, 20^. In reality this muscle contraction is a combination of both monosynaptic and polysynaptic connections mediated via premotor neurons, that converge on the motor neuron. The Hoffman (H) reflex is an artificially induced counterpart whereby Ia afferents are stimulated electrically and the resultant compound EMG in a distal muscle is measured. This has the benefit of being minimally invasive and removes variation from muscle spindle activation. The proprioceptive reflex circuit is open to modulation from numerous sources such as inhibition from antagonistic muscles, Renshaw inhibition and effects of descending motor pathways ^21, 22^. Under normal physiological conditions repeated firing of muscle spindles, such as during voluntary movement, causes a near full suppression of the reflex, termed rate dependent depression. This response is attenuated or lost in many SCI patients such that the reflex remains large ^16, 23^.

The exact mechanism by which spasticity and hyperreflexia develops is unknown. A recent study in shows that spared reticulospinal function is positively correlated with the presence of spasticity in SCI patients ^24^. In animal models a number of anatomical and molecular changes associated with hyperreflexia have been reported; alteration in the chloride ion transporter KCC2 in motor neurons ^25^, persistent inward currents driving increased motor neuron excitability ^26^, increased excitatory input from Ia afferents ^5, 27^ and reduced inhibition on the circuitry ^28^. Whilst the symptoms of spasticity are amenable to pharmacological agents like Baclofen, a centrally acting GABAB receptor agonist, or α-2 adrenoreceptor agonists like tizanidine, these agents are associated with numerous contraindications and side effects at higher doses ^29^. Currently no treatment exist that correct the underlying pathology.

NT3 is an attractive treatment option based on its role in muscle afferent development. The survival and differentiation of Ia afferent neurons during development is dependent on neurotrophin-3 (NT3). A pro-form is synthesised and cleaved into a mature 15kDa protein which has been shown to be constitutively secreted from cultured neurons, however the pro-form also has the capacity to be released ^30–33^. Either form is suspected of being secreted from muscle spindles where it acts as a guidance molecule for the incoming afferents. NT3 is essential for the correct wiring of Ia afferents: without it the axons fail to innervate the peripheral muscle spindle and/or fail to project deep into the ventral horn to reach their central targets ^34^. Beyond development, NT3 regulates the strength of sensory-motor monosynaptic connections in both intact and axotomized stretch reflex arc circuits ^35, 36^. Additionally, NT3 is suspected to play a role in the normal functioning of descending pathways involved in locomotion, such as the corticospinal tract ^37^ and serotonergic fibres ^38^, which continue to express the high affinity receptor in adulthood.

NT3 binds primarily to tropomyosin kinase receptor type C (TrkC), its high affinity receptor, however it has the capability to bind and signal through the other Trk receptors ^39^. It can also signal through the promiscuous neurotrophin receptor p75, which is involved in generating both pro-survival and pro-apoptotic states ^40^. Whilst NT3 levels significantly drop in the muscle beyond the postnatal period ^41^, the expression of its receptor TrkC remains high in large diameter dorsal root ganglia (DRG) neurons ^42, 43^ indicating mature proprioceptive neurons remain receptive to NT3.

Previous studies highlight NT3 as a potent inducer of neuronal plasticity following injury. After exposure to increased NT3 at the lesion site, CST axons which have been transected undergo increased axonal sprouting ^44^. After unilateral pyramidotomy, the intact CST responds in a similar way in the presence of elevated (transgenic) NT3 in the lumbar cord with enhanced sprouting across the spinal midline towards the ipsilesional grey matter ^45, 46^. An increased level of NT3 is known to improve hindlimb function after; retrograde transport of NT3-expressing vectors to motor neurons ^47, 48^, direct delivery of NT3 to the lesion site through intraspinal gene therapy ^49^, intrathecal injection ^50^ or via NT3 secreting fibroblasts ^51^. Rostrally located fibres which are spared after a partial CST injury are also responsive to elevated NT3 ^52^. These functional effects could also be a result of NT3 exposure strengthening monosynaptic proprioceptive afferent connections within the ventral horn ^35^. Additionally, endogenous levels of NT3 in the spinal cord can be raised following exercise in intact and spinally transected rodents ^53–55^.

Fewer studies have focussed around whether NT3 can improve function in the forelimb following SCI, with those performed finding differing effects depending on location of the injury and NT3 treatment. Overexpression of NT3 in the reticular nucleus, the origin of reticulospinal tract fibres did not enhance pellet reaching recovery above control treated animals ^56^. Following CST injury in the brainstem, overexpression of NT3 in forelimb flexor muscles and raised serum levels were able to partially restore several sensorimotor forelimb tasks, which interestingly coincided with complete reduction in hyperreflexia ^57^. Pre-treatment of muscles four weeks prior to injury with an adeno-associated viral vector (AAV) encoding NT3 was able to partially overcome forelimb deficits caused by dorsal column transection at the cervical level ^58^. A combined therapy of NT3 and the anti-inflammatory cytokine interleukin-10 loaded onto a biocompatible nerve bridge has recently been shown to convey functional benefits in the affected forelimb after cervical lateral hemisection ^59^. Treatment was delivered at the time of injury and shows synergistic effects can be achieved when combining NT3 with a pro-regenerative cellular environment.

### Rationale for testing Intramuscular NT3 in a cervical contusion

To date no one has assessed the therapeutic effects of NT3 as a monotherapy after cervical contusion. For patients with the highest grade of SCI that impairs function in both upper and lower limbs, the restoration of useful hand function is their main priority, where it would allow for greater independence ^60^. Hyperreflexia develops in the forelimb in rodents after a moderate mid cervical contusion injury (Sydney-Smith et al., unpublished observations). Here, we set out to test the hypothesis that neurotrophin 3 would normalise the deficits in sensorimotor function, exaggerated spinal cord reflexes and changes to spinal cord connectivity that follow from cervical SCI. To maintain clinical relevance, the mid cervical injury was used again to model the most frequent location of injury in humans. Subcutaneous delivery of NT3 protein has been proven safe and well tolerated in patients with other conditions ^61^, and an AAV encoding NT3 gene therapy approach is the focus of an ongoing Phase I/IIa safety and efficacy trial for Charcot Marie Tooth 1a neuropathy (NCT03520751). Through the use of intramuscular viral delivery of NT3 in this study it was hoped to generate a continual elevation of NT3 within the muscles of the forelimb in addition to raised levels of in the circulation.

The exact mechanism by which the intramuscularly overexpressed NT3 conveys a therapeutic effect after SCI is unknown. This pool of NT3 protein in the muscles is assumed to bind to proprioceptive afferents, which express TrkC, and be trafficked to the DRG where it can induce transcriptional changes and possibly alter the connectivity and function of these afferents ^62, 63^. NT3 has the capacity to be retrogradely transported in motor neurons, providing an additional route to the proprioceptive circuitry ^64^. As NT3 can cross the blood- nervous system barrier, any elevation in serum could have an effect along the neuraxis ^46^. The DRG might be particularly amenable to therapeutic effects through raised serum NT3 due to its dense fenestrated vasculature ^65, 66^. In summary, delivery within the muscle offers a clinically applicable delivery route whilst enabling it to be trafficked to different neuronal targets, and possibly act through both sensory and motor components of spinal circuits as well as systemically across the blood-CNS barrier.

## Methods

### Animals

All animal procedures were conducted in accordance with the UK Home Office guidelines and the Animals (Scientific Procedures) Act of 1986. A total of 30 Lister Hooded rats (Charles River), weighing 240-270g were used at the start of the study, with five animals being excluded from the study prior to surgery due to poor reaching ability following training and a further three dying due to anaesthetic complications at varying stages of the study. Animals were group housed in standard conditions with a 12 hr light/dark cycle and access to chow food and water *ad libitum.* Animals were acclimatised in the unit for at least one week prior to any behavioural or surgical procedures but during this time animals were handled daily. To acclimatise them to the post injury recovery diet animals were offered high calorific food such as; peanut butter, chocolate spread and seeds, in addition to nutritional and hydration supplements (Dietgel #76A and Hydrogel, ClearH_2_0) approximately one week prior to injury.

### Experimental Design

The study was a stratified block design. Animals were ranked based on their final session of single pellet reaching during baseline sessions and allocated to 3 groups; contusion + NT3 (n=12), contusion + PBS (n=12) and uninjured + no treatment (n=5). This was achieved by allocating the best performer into the first treatment group, the next best into the second treatment group, the third best into the first treatment group and so forth. Every fifth animal was allocated as a naïve animal (**Figure 1**).

**Figure 1.**
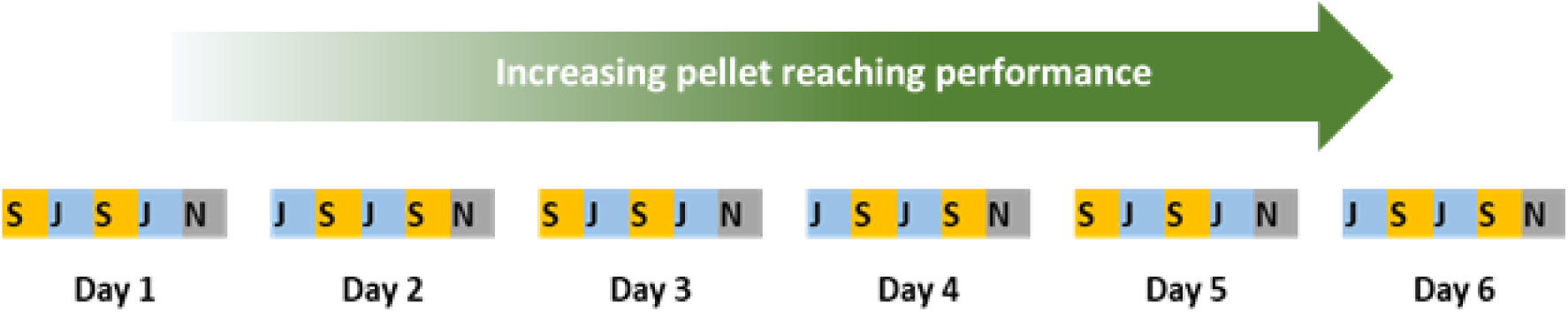
Allocation of animals into treatment groups based on pellet reaching performance. Animals were scored prior to injury, ranked from worst to best and sequentially allocated into one of two treatment groups. Every fifth animal was designated a naive animal. Surgeries began with the worst performers and on any single day there was an equal number from both treatment groups being operated and treated. S&J refer to the blinded treatments of either NT3 or PBS. N refers to naïve animals which did not undergo contusion injury or intramuscular injection.

This way each group would have an even spread of good and poorer performers on the pellet reaching task.

Randomisation occurred at the level of the group: each contusion group was randomly allocated to receive either PBS or NT3. An independent person (Zoe Hore) coded the vial of AAV-NT3 and the vial of PBS as “J” or “S” so that all surgeons, observers, analysts and PI were blinded to treatment allocation. Blind-coded single-use aliquots were used and codes were only broken at the end of the experiment. Surgeries were performed during a single week starting with the poor pellet reaching performers and on each day there was an even spread of animals from both treatment groups. Because of this, animals were housed in mixed cages comprising injured animals of both NT3 and PBS treated animals. Naïve animals were housed together in a separate cage. The order of surgeries within a day was alternated, so that each day began with a different group (Figure 1). Each experimental animal had a Radio-frequency identification microchip inserted subcutaneously and this was used to identify each animal throughout the study. Blinding was broken once the ELISAs were performed; this occurred prior to horizontal ladder analysis but after all other behavioural and H reflex data was analysed. Only once the horizontal ladder data was assessed were the individual microchip numbers matched back up to the two treatment groups of ‘J’ or ‘S’, and then designated NT3 or PBS treated groups.

### Cervical contusion injury

Rats were anaesthetised with Isoflurane, 5% for induction, reduced to 2% for maintenance during surgery, with oxygen flowing at 1L/min. Preoperatively 5mg/kg Enrofloxacin (Baytril) and 5 mg/kg carprofen (Carprieve) were injected subcutaneously. The region around the cervical and upper thoracic neck was shaved, the skin prepared with both 4% w/v chlorhexidine gluconate (Hibiscrub) and iodine antiseptic (Vetark Professionals). Finally, eye ointment (Viscotears) was applied. The animal was positioned prone on a homeothermic blanket (Harvard Apparatus), a rectal thermometer inserted to maintain a body temperature around 36°C.

A 2 cm incision was made through the skin, from below the occiput to the T2 vertebral prominence to expose the underlying neck muscles. The hibernation gland and fat tissue between the scapulae were reflected caudally and the muscles overlaying the middle cervical vertebrae divided in separate layers. Using the spinous process of T2 as a landmark, and noting that rodents have seven cervical vertebrae, the lamina belonging to the C5 and C6 vertebrae were located and attached muscle removed. A bilateral laminectomy involving the entire C6 lamina and the caudal half of C5 was performed, exposing approximately 2mm either side of the midline. Loose cotton swabs were used to stop any bleeding and any sharp bone fragments within and around the laminectomy were carefully removed. The dura remained intact. The animal was transferred to the Infinite Horizons impactor (Precision Systems Instrumentation) and secured in place by two Addison forceps clamped on to the lateral processes of C5 and C7 vertebrae. A bilateral contusion, 225 kDyn with 0s dwell time was delivered using a 3mm diameter custom made tip (Precision Systems Instrumentation), corresponding to the region between the C6 and C7 spinal segments. Impact force and tissue displacement was recorded, and no impact produced a force/displacement curve suggestive of hitting bone or the animal slipping within the device. Sham animals (n=5) in this study did not received any surgery nor post-operative pain medication and are subsequently referred to as naives.

The contusion site was inspected for bleeding and the muscles sutured in three layers using 4-0 Vicryl absorbable sutures (Ethicon). All animals received 5ml saline subcutaneously and placed in a 30°C water-based incubator (Thermocare) whilst the anaesthesia wore off. For at least the first four days after injury animals continued to receive; warm saline subcutaneously to combat dehydration, 5 mg/kg Baytril for potential bladder infection and 5mg/kg Carprieve for pain control. Rigid paralysis occurs in the forelimbs after injury and there is minimal hindlimb activity for the first few days post injury. During this time animals were laterally recumbent, which necessitated housing on soft paper bedding and being handfed high calorific food and water every few hours. Food and hydrogel were located around the cage and within reaching distance of their mouths. By one week after injury all animals regained the ability to weight-bear and reach food and were placed back onto sawdust in cages in the original holding rooms. Animals were able to void their bladders adequately and whilst bladder volume was checked during post-operative care, no manual expression was necessary.

### Adeno-Associated Viral vector encoding Neurotrophin 3

An AAV transfer plasmid, AAVspNT3, contained human prepro-Neurotrophin 3 under a CMV promoter. The neurotrophin-3 coding DNA sequence (NM_002527.4), corresponding to isoform 2 precursor protein including the secretory signal, was flanked by splice donor/splice acceptor sites and terminated in a beta globin poly(A) sequence, a gift of Prof. Fred Gage, Salk Institute. This was packaged into an adeno-associated viral vector serotype 1 by the University of Pennsylvania Viral Vector Core facility (custom batch CS1232). This viral vector is referred to in the text as AAV-NT3. The viral vector titre was measured by the Viral Vector Core using digital droplet PCR (6.2 x10^13^ Genomic Copies (GC)/ml) and in our lab via qPCR (1.22 x10^13^ GC/ml) by Dr Aline Spejo. The latter value was used for calculations when diluting AAV-NT3. Primers for the qPCR were: Forward 5’-AATTACCAGAGCACCCTGCC and Reverse 5’- TTTTGATCTCCCCCAGCACC.

### Intramuscular injections

Twenty-four hours after the contusion surgery the muscles of the dominant forelimb, previously determined from single pellet reaching performance, were injected with a total of 1.04x10^11^ GC of AAV-NT3 diluted in 160 μl of PBS. Glass Hamilton syringes attached to a 26G metal needle with a non-coring bevel were used throughout with care taken to insert the needle bevel facing up and distal to the final injection site. Injections were either superficial, where the bevel of the needle was still visible through the fascia, or deep, where the bevel was no longer visible and resided in the remaining muscle bulk (**Figure 1**).

Animals were anaesthetised with isoflurane and given eye ointment as before. Enrofloxacin and Carprieve was already given as part of the early morning post-operative care following the contusion injury, however an additional half dose (2.5mg/kg) of Carprieve was administered to animals that underwent intramuscular injections in the afternoon. The dominant forelimb was shaved and the skin and paw prepared as before. For all the muscle injections, the muscles were injected through the fascia and apart from the abductor digiti minimi (ADM), all muscle injections were made parallel to the bone. Following each set of muscle injections, the incision was sutured with 4-0 Vicryl sutures before moving onto the next. Muscle injections occurred in the following order: flexor compartment, extensor compartment, biceps brachii, triceps brachii and then ADM.

For the forelimb flexors, the animal was positioned supine and the forearm was extended. A 1cm incision was made through the skin just below the elbow and the skin was retracted with blunted hook retractors. The flexor muscle compartment was injected via a series of six deep injections of 5ul each, running in a circumferential arc across the palpable fascial compartment. The needle was inserted close to the wrist and ran proximally whilst keeping parallel to the bone. The bevel was inserted two thirds of the distance proximodistally from wrist to the elbow before injecting. For forelimb extensors, the dominant forearm was supinated and taped above the animal’s head. A similar incision was made as before on the radial side and skin was retracted. The extensor compartment was injected in a similar fashion with six deep injections of 5μl each in an arc approximately two thirds up for distal forearm.

For Biceps Brachii, the arm was extended and supinated. A 1cm incision was made above the antecubital fossa and the skin was retracted. A total of three deep and three superficial injections, each 5 μl, was made, targeting approximately halfway between the insertion and origin of the biceps muscle. For Triceps Brachii the animal was positioned similarly for the extensor compartment and a slightly longer incision made from above the elbow proximally towards the shoulder. The skin was blunt-dissected and the long head of the triceps was exposed. This was injected with five deep injections of 5μl each and three superficial injections of 2.5μl each. The forearm was then further extended to expose the lateral head of the triceps from the same incision. This was injected with five deep injections of 5μl and three superficial injections of 2.5μl. Finally, the ADM muscle was injected through the paw skin with one deep and one superficial injection of 2.5μl each.

### Single Pellet Reaching

Ability to successfully reach, grasp and return to eat 45mg pellets was assessed every two weeks in modified Whishaw window devices. Rats were placed in a Perspex cage with a 15 mm wide slit to allow for reaching. The sugar pellets were presented on a ledge 15 mm away from the reaching window and 5mm lateral from the edge of this window (**Figure 9A**). This gap limits grasping that uses compensatory reaching mechanisms such as dragging pellets.

Animals were pretrained for several weeks prior to the study. Initially sugar pellets were positioned in the centre to facilitate learning and to determine the dominant paw. A sugar pellet was placed only once the animal’s forepaw had both touched the ground and had repositioned. Animals were subsequently trained every other day for three weeks, with each session comprising 10 minutes of pellets being offered to the dominant paw. When scoring sessions, baseline or after injury, an initial ten pellets were given as a warm-up (which were not scored) followed by twenty pellets which were scored using the following definitions:

1. Success on first attempt - Animal retrieved and ate the pellet in a single continuous motion.
2. Success with multiple attempts – The animal retrieved and ate the pellet but took two or more attempts to do so. If the animal made five attempts and then retrieved the hand through the window the trial was recorded as a miss. This was rare as animals would usually knock the pellet off the ledge during the initial few attempts.
3. Miss- either the animal failed to reach the pellet within five attempts, or the pellet was knocked off the platform or it was dropped within the cage prior to being eaten.

Baseline scores were averaged from two separate behaviour sessions. Owing to the low percentage of ‘success on first attempt’ being achieved at the end of training, animals were ranked according to percentage of ‘Success with multiple attempts’ before allocation into surgical groups.

### Horizontal ladder

Ability to accurately place the forepaws during locomotion was assessed with the horizontal ladder every two weeks. Rats were filmed crossing a horizontal ladder with one metre of randomly spaced rungs ranging from 1-3cm apart. Each session, a total of three complete runs were filmed per rat and any run where the animal stopped for more than one second was discarded. Videos were analysed and steps on both forelimb sides assessed using seven point horizontal ladder scoring system ^67^. Briefly, a score from zero to two indicated slip based errors; zero is a total miss where the forelimb did not contact the rung at all, one is a deep slip where the limb weight bears but slips causing disrupted locomotion and two indicated a slip similar to previously but without interruption to gait cycle. Scores of three to five represent placement errors; three indicates a step whereby the animal initially places the paw but then replaces it on another rung prior to weight bearing, four indicates the paw was either adjusted on the same rung or was repositioned to another rung prior to contact and finally five indicates steps where the wrist or a few digits weight supported the animal as the paw did not make full contact with the rung. A score of six indicates an error free step. The initial and final step were omitted from analysis and only steps that fell on the irregular spaced rungs were assessed.

### Grip strength

Grip strength was assessed every two weeks using a Dunnett Style Grip Strength Meter (Linton Instrumentation). This device comprises two collinear bars attached to separate force transducers and is capable of recording grip strength of each forelimb independently yet simultaneously. Rats were held around their hip and were lowered over the device until rats grabbed both bars with all forepaw digits. Steady but firm pressure was used to pull horizontally on the base of the rat’s tail until it released its grip. Measurements were averaged over three trials taken within the same testing session. This task was performed last to avoid any negative effects from the stressful exertion on the performance in other tasks.

### Retrograde tracing

Cholera Toxin Subunit B (#104, List Biologicals) was used to retrogradely trace motor neurons of the distal forearm three to four days prior to perfusion. Animals were anaesthetised and both arms prepared for surgery as before for muscle injections. An incision was made on the ulnar side of the distal forearm and then skin was retracted. The fascia covering the Flexor Digitorium Profundus and Flexor Carpi Ulnaris was cut and blunt dissection used to free up the ulnar nerve where it bifurcates distal to the elbow. Using a 34G needle on a Hamilton syringe, a total of 1μl of 1% Cholera Toxin B diluted in ddH_2_0 was injected slowly into both ulnar fascicles. The fascia and skin were sutured. Animals did not appear to suffer any loss of function of the distal forearms due to the injections, however no behaviour was assessed after tracing. One animal from the NT3 group died during anaesthesia.

### Tissue collection

Animals were anaesthetized with 80mg/kg of sodium pentobarbital (Euthatal) and the thoracic cavity opened. Approximately 1.5ml of blood was taken through cardiac puncture of the left ventricle and stored on ice. The animal was then transcardially perfused with 80ml ice cold PBS and the distal half of the biceps brachii muscle of the dominant limb removed and snap frozen in liquid nitrogen. Animals were then transcardially perfused with 4% paraformaldehyde diluted in PBS over the course of ten minutes. Inspection of the vertebral column during dissections confirmed that the impact site was between the C5 and C6 vertebral lamina in all animals. The cervical spinal cord from segment C3 – T2 and the C4-T1 dorsal root ganglia (DRGs) were post fixed in fresh 4% PFA for 24 h at 4°C. Whole blood was kept overnight at 4° and snap frozen tissue kept at -80 °C

### Immunohistochemistry

Spinal cord and DRGs were removed from PFA, washed 3x5 mins in PBS and stored in PBS with 0.02% sodium azide at 4°C. Cervical dorsal root ganglia were transferred to 30% sucrose with 0.02% sodium azide in PBS for a minimum of two days to cryoprotect prior to histological analysis. Following MRI, spinal cords were washed several times in PBS before being cryoprotected as above for a minimum of three days until the tissue sunk. The region caudal to the epicentre cavity was then mounted and frozen in Optimum Cutting Temperature Medium (VWR International), cryosectioned in series at 20μm thickness onto Superfrost slides (Fisher Scientific Ltd) and stored at -20°C

Prior to immunohistochemistry the sections were baked for 30 mins at 50°C and residual OCT removed in 2 x 1min washes in PBS. Sections were blocked in 3% BSA (#A3059, Sigma) in 0.05% Triton-X-100 (#T8787, Sigma) in PBS for 1h at room temperature. This was then tipped off and the slides incubated in the relevant primary antibody (Table 1) diluted in 0.05% Triton-X-100 in PBS with 1% BSA, overnight in a humidified chamber at 4°C. Sections were then washed in 3x5 mins washes of PBS and incubated with the relevant secondary antibody (Table 1) in 0.05% Triton for two hours at room temperature. Sections were finally washed, 3x5 mins in PBS, mounted with Fluoromount-G Mounting Medium with DAPI (Invitrogen). Care was taken to ensure sections never dried out during the staining.

**Table 1:** List of primary and secondary antibodies.

| Antibody | Dilution | Supplier |
| --- | --- | --- |
| **Rabbit-anti-Vglut1** | 1:1000 | #135 302 Synaptic Systems |
| **Goat-anti-CTB** | 1:2000 | #703 List Biologicals |
| **Mouse-anti-NeuN** | 1:500 | #MAB377 Millipore |
| **Goat-anti- mouse Dylight 650** | 1:500 | #AB96882 Abcam |
| **Donkey-anti- goat Alexa 546** | 1:1000 | #A-11056 Invitrogen |
| **Donkey-anti- rabbit Alexa 488** | 1:1000 | #A21206 Invitrogen |

### Quantification of vGLUT1+ immunoreactivity

A total of four to five sections, evenly spaced throughout the C8-T1 segments, were imaged on an LSM 710 inverted confocal microscope (Zeiss). Successfully traced α-motor neurons (Ctb+/ NeuN+) in the ventral horns were taken at 400x magnification, with aperture and exposure settings for each of the four channels kept consistent throughout. Images were analysed in FIJI. A region of interest (ROI) was drawn around NeuN+ motor neurons with a clear nucleus using the polygonal selection tool. The area and perimeter were measured using the Ferets Diameter function. Then this ROI was expanded into a band 2.5μm wide using the ‘make band’ function and the background subtracted from the vGlut1 channel using the rolling ball function, radius =10. The integrated density was measured, which gives the raw integrated density. This allocates a value for each pixel on a scale of 0-255 in which 0=black/no signal and 255 = maximum brightness. Finally, each puncta that could be resolved individually within this 2.5μm band was counted and reported as a vGlut1+ apposition. An overview of this analysis is provided in (Figure 16)

### ELISA & BSA

Levels of NT3 protein in serum and muscles were assessed using a Human Neurotrophin-3 ELISA Kit (#ab100615 Abcam). In this study the viral vector encoded the human prepro-NT3 gene, which when cleaved from their pro-domains, is identical to mature rat NT3. This means any endogenous NT3 is also detected by this assay. The lower limit of detection is 4.12 pg/ml. Whole blood was centrifuged at 10,000 rpm (8200 g) at room temperature, and the serum removed without agitating the clot. Serum was spun again for a further 5 mins, aliquoted and stored at -80°C until used. Muscle samples were weighed and 10ml per 1g of tissue in Tissue extraction Reagent I (#FNN0071 Invitrogen) with added protease inhibitor (#S8820 Sigma-Aldrich) was added to Gentle MACS M tubes (# 130-096-335 Miltenyi Biotec). The tissue was kept on liquid nitrogen until ready to digest, when samples were placed in the buffer and immediately homogenised using a Gentle MACS Dissociator (Miltenyi Biotec). Homogenates were then centrifuged at 4000rpm (2808 g) for 10 mins at 4°C. The supernatant was aliquoted and centrifuged again at 4°C and stored at -80°C until used.

ELISA was performed according to manufacturer’s protocol. Dilutions were previously determined such that NT3 levels fall within the standard curve. Briefly, serum was allowed to thaw on ice before being diluted 1:2 in Diluent A. Muscle extractions were diluted 1:250 for NT3 treated group and 1:6 for the uninjured and PBS groups in Diluent B. The standards and treatment samples were prepared and run in triplicate. Samples were incubated on the plate for 2.5h at room temperature, washed in supplied wash solution and incubated for 1h at room temperature with biotinylated NT3 antibody. This was washed off and HRP-streptavidin applied for 45mins, before being washed off. The substrate reagent was added for 30mins in the dark, followed by the stop solution and the plate read immediately at 450nm using a BMG LabTech FLUOStar (Omega). NT3 levels were normalised to the total amount of protein extracted from each sample.

Alongside the ELISA a Bicinchoninic acid assay (BCA) assay (#71285-3, Novagen, Millipore) was performed according to manufacturer’s protocol. Serum samples were diluted 1:150 in PBS, and muscle homogenates were diluted 1:20 in PBS. A four parameter standard curve was generated for both assays and was above r>0.99

### Ex vivo MRI

For lesion quantification the epicentre of the lesion was imaged using a 9.4 T MRI scanner (Brucker Biospec). Spinal cords were wrapped in loosened 8-ply surgical cotton gauze swabs ( Premier Healthcare Excellence) and inserted into a 50ml Falcon tube containing a custom made 3D-printed barrel to immobilise the spinal cords . The tissue was fully submerged in Fomblin (Solvay), a proton free fluid which reduces background signal, and all air bubbles were removed prior to scanning. A T2 weighted image was acquired using a fast spin-echo sequence and the following parameters: echo train length = 4, effective TE = 38 ms, TR = 3000 ms, FOV = 40 x 20 x 20 mm, acquisition matrix = 400 x 200 x 200, acquisition time = 9 h 20m. This composite scan contained T2W images from 14 spinal cords scanned simultaneously.

ITK -snap and Convert3D software was used to process the T2 weighted composite. Using Convert 3D and the ‘region’ command, a 5x5x5mm region of interest (ROI) comprising the epicentre was selected for each individual spinal cord. This was the extent of the observable lesion, defined as the region where the first changes in the spinal cord architecture were appreciated on the transverse view and most often was a loss of grey/white matter distinction in the dorsal funiculus. The ROI was resliced using the ‘resample’ command of Convert 3D to generate a scan comprising 100 slices with an isotropic voxel size of 50μm. Within ITK-snap the contrast was increased and thresholding, lower limit=0.110 & upper limit=0.312, applied to select white matter (plus any other tissue with equivalent MRI signal). The automatic segmentation tool was initiated on the thresholded image at 2 sites of intact white matter, one at each of the rostral and caudal most slices and left to run until both initiation points joined. The segmentation parameters were as follows: competition force =1.000, smoothing force = 0.05, α=1.000, β=0.100, speed = 5.00 initially but reduced to 1.00 once segmentation was complete to smooth the mask. All parameters of contrast and thresholding were identical for each spinal cord. The mask was exported as a screenshot series of the transverse view and images every 150μm were quantified in FIJI. For spared white matter area a custom macro was generated which; converted the series to a stack, isolated the red channel, used the ‘create selection’ tool to select the red region corresponding to the segmented ROI and measured the total area for each slice. Volume for each parameter was calculated as the thickness of the voxel multiplied by the cross-sectional area. For total spinal cord volume, the additional step ‘Analyse particles including holes’ was added, before creating the selection.

### H reflex recording

Rats were anaesthetised with an intraperitoneal injection of 0.10 mg/kg medetomidine and 30mg/kg ketamine diluted in saline. The correct plane of anaesthesia for testing was indicated by the absence of whisker movement and absence of hind paw pinch reflex. Rats were positioned supine and a rectal temperature probe inserted and connected to a homeothermic blanket (Harvard Apparatus) to maintain a constant body temperature of approximately 36°C. Eye ointment (Viscotears) was applied and the wrist and forepaw cleaned with 4 % chlorhexidine. The forepaw digits were taped down such the 4^th^ digit was not ad/abducted. Two 26-gauge needles were positioned subcutaneously in parallel and approximately 2mm apart, over the carpal bones and connected to stimulating electrodes. A single 30-gauge concentric needle recording electrode (Digitimer) was positioned superficially into the ulnar aspect of the forepaw, no more than 1-2mm deep, to record compound muscle action potentials (CMAPs) of the ADM muscle. Minor alterations to the positions of recording and/or stimulating electrodes were made to obtain maximum quality CMAPs prior to recordings.

A monophasic square wave stimulus, of width 100µs, was applied via an isolated constant current pulse stimulator, (NL800; Digitimer). The CMAP signal was passed through a differential amplifier, amplified 2000-fold and run through a band-pass filter that retains signal between 300Hz – 6kHz to remove interference originating from mains electricity. The signal was then digitized via PowerLab and visualised in LabChart. Threshold was defined as the stimulus in which the M wave was elicited in three out of four stimuli. A recruitment curve was generated using stimuli starting at 1x motor threshold (MT) to a maximum of 2x MT in 10% increments. Rate dependent depression (RDD) was tested at 1.2x MT by first delivering 25 pulses of 100µs width at 0.2Hz, and 30s later delivering a train of 25 pulses, at 5Hz (**Figure 3A**)^68^. Recordings were repeated on the opposite forelimb, with an additional half dose of anaesthetic given if 30 mins had elapsed. If so, a 10 min period was allowed for the animal to return to a stable plane of anaesthesia. Once recordings were finished, 2mg/kg atipamezole hydrochloride intraperitoneally was used to reverse the medetomidine and 5ml saline subcutaneously to counter dehydration during recovery. Animals remained on a heat blanket until upright and moving and returned to an incubator overnight. The likely current delivered to the animals was determined empirically by measuring the voltage across a 1k Ohm resistor that was connected to the outputs of the constant current generator. Both treatment groups were tested at 1, 3, 5, 7, 9, 11, 13 weeks post injury. Uninjured animals were tested at 0, 3, 7, 11, 13 weeks post injury

**Figure 3.**
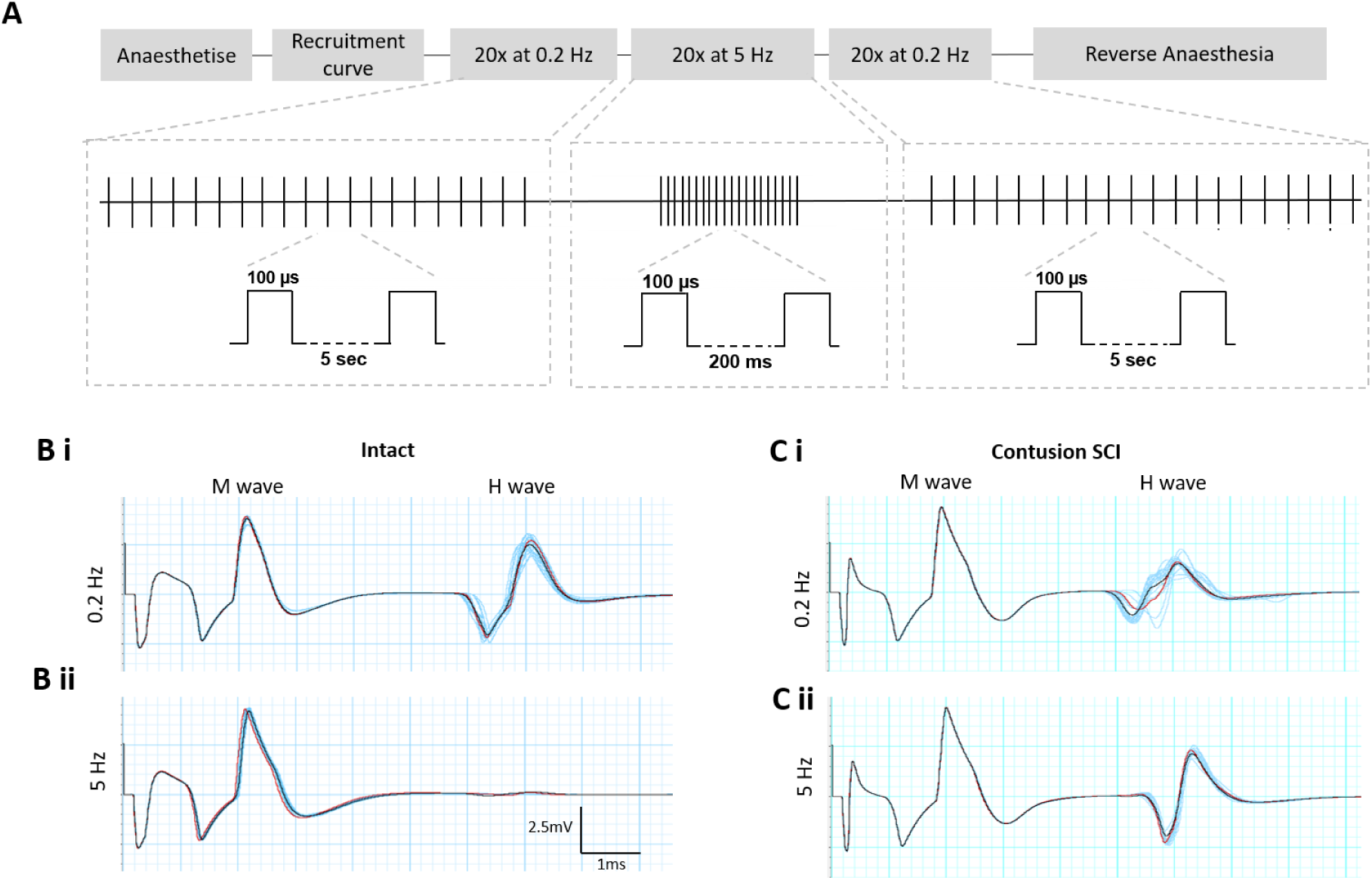
H reflex stimulation protocol to assess hyperreflexia in forelimb following SCI. **A)** Following acquisition of a recruitment curve each animal was assessed for hyperreflexia using a train of electrical stimuli. Initially 20 stimuli at a low frequency of 0.2Hz was delivered, followed by a train of 20 stimuli at a higher frequency of 5Hz to elicit rate dependent depression of the H reflex. Afterwards the stimuli at the lower frequency were repeated for control purposes. A monopolar stimulus of 100μs duration was used throughout. **B&C)** Representative traces of an animal before, and five weeks after spinal cord contusion. All 20 traces are shown in blue, with the average overlaid in black and last trace in red. **Bi)** At the slower frequency both a Motor and H wave are present. **Bii)** However, during the train of stimuli at higher frequency the H wave is fully depressed. **Ci** & **Cii)** Following injury, the H wave fails to depress and remains large.

### H reflex analysis

The absolute amplitude for each M and H wave was measured in Labchart and averaged for each stimulation intensity. From this the maximum size of H wave (Hmax), and then maximum size of M wave (Mmax) were identified and the Hmax:Mmax calculated. To assess RDD, the amplitude of the average H wave at either 5s or 0.2s interstimulus interval was normalised to the Mmax for that limb during the testing session. RDD was then defined as the amplitude of the normalised H wave at 0.2s interval as a proportion of the H wave at 5s interval and represented as a percentage.

### Statistical analysis

Statistical analysis was performed in either Prism 8 (GraphPad) or SPSS (IBM) software packages. Behavioural and electrophysiology data were analysed using a linear model with a covariate structure best suited to the data being analysed ^69^. The use of restricted maximum likelihood allows for unbiased estimation of model parameters taking into account different variance and covariance structures. It also enables data sets with missing values to be analysed without the exclusion of entire animals from this longitudinal study. Three models were generated using either: Unstructured, Compound Symmetrical or First-order autoregressive covariate structures and analysis performed on the model with lowest Akaike information criterion, a measure of model fit. Baseline measurements were included as a covariate in the model unless stated in figure caption.

For complete data sets the following tests were performed where appropriate in Prism8 (GraphPad); Paired Student T-tests, one way & two- way ANOVAs. The tests performed are stated for each associated figure, along with number of subjects, p value and F statistic for ANOVAs. Error bars are mean ± Standard Error of Mean.

## Results

### Overview

A moderate contusive injury of 225kdyn was delivered to 24 Lister Hooded rats between the C5 and C6 laminae, corresponding to the C6-C7 spinal level. Contused rats either received 1x10^11^ GC of AAV-NT3 in 165μl PBS (termed NT3 group, n=12) or an equivalent volume of PBS (termed PBS group, n=12). Uninjured rats served as the injury control for this study and received no injury (termed naïve group, n=5). Starting one week after injury the degree of forelimb hyperreflexia was assessed every other week until the end of the study. Assessment of sensorimotor function took place at two weeks post injury and then every subsequent two weeks (Figure 2A). At the end of the study afferents and motor neurons were retrogradely labelled with CtB injections into the ulnar nerve bilaterally; tissues and blood were collected for histology and ELISA, respectively, after four days. In order to maximise the level of NT3 available to the sensorimotor circuitry below the level of the injury the majority of forearm muscles were injected in the dominant limb (Figure 2B). An injection protocol was devised which targeted the motor end plate regions ^70^. Since three-dimensional imaging has revealed motor plates can lie deep within the muscle ^71^, this protocol involved both superficial and deep injections in a circumferential arc around the muscle belly (Figure 2C & D). This protocol results in widespread transduction of myocytes throughout the main bulk of the muscle as shown in a separate experiment, using AAV1-GFP under the same promoter, within the lab.

**Figure 2.**
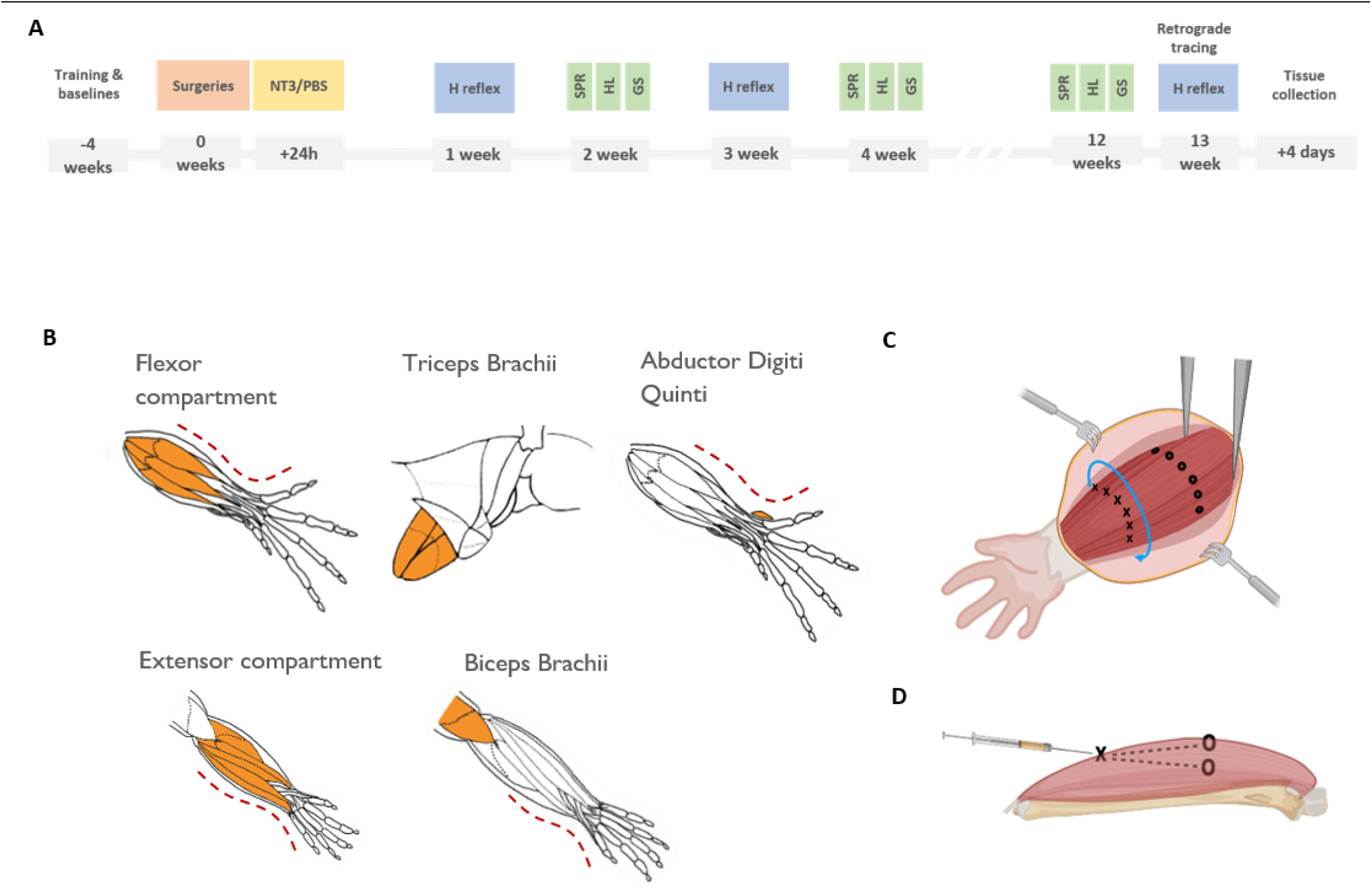
Experimental overview. A Experimental timeline, SPR= single pellet reaching, HL = horizontal ladder and GS=grip strength. B Schematic of the right forelimb, in supination at top and pronation on bottom, showing the muscles or compartments which were injected with either NT3 or PBS. For flexor and extensor compartments individual muscles are shown and these were located approximately through the fascia. For reference, ulnar side is marked with a thin red line. C & D The muscle injection procedure. C Individual muscles or the compartments beneath the fascia were emphasised using forceps. Injections were performed in a circumferential arc around the muscle belly, shown in blue. Each compartment was divided into 3 along the proximodistal axis, with the location of needle insertion shown by X, and O indicating the final location of the bevel. D Cross section through a muscle indicating the approximate location of both the superficial and deep injections. Both injections had similar location of insertion and ran parallel to the long bone. Whilst the bevel was visible during superficial injections, this was not the case for deep injections as the bevel resided within the muscle belly. Fig 2B adapted from ^70^

### NT3 treatment was unable to correct hyperreflexia in distal forelimb circuitry

To measure hyperreflexia after injury, H reflex recordings were performed on both injected and noninjected forelimbs before surgery and then every two weeks in the NT3 and PBS group and every four weeks in naives. Previously it has been shown that stimulating the ulnar nerve at a frequency of 5 Hz during paired pulses is sufficient to induce rate dependent depression in the ADM muscle ^57^. It was thought that stimulating the nerves at the wrist with a train of stimuli at 5Hz frequency would drive as much depression of the H wave as possible (Figure 3A). This technique has been used previously in the rodent forelimb after spinal cord injury^68^. Stimulating the wrist at 1.2x motor threshold intensity with this frequency and recording compound motor action potentials (CMAPs) in the ADM muscle generated distinct motor (M wave) and H reflex (H wave) responses (Figure 3Bi). In intact animals the H wave diminishes during this higher frequency train indicating rate dependent depression (compare Figure 3Bi with Figure 3Bii), whilst in injured animals the H reflex remains present (compare Figure 3Ci with Figure 3Cii). The amplitudes of the M waves were not affected by this frequency of stimulation, as expected ^57, 68^.

Prior to injury all animals exhibited rate dependent depression of the ADM H-reflex circuitry (Figure 4). On average the mean H wave amplitude was depressed by about 92-95% during the higher frequency stimulation compared to the size of the H wave when stimulated at the slower frequency. Before injury, all three groups showed rate dependent depression in both the dominant and nondominant paws, as expected (Figure 4).

**Figure 4.**
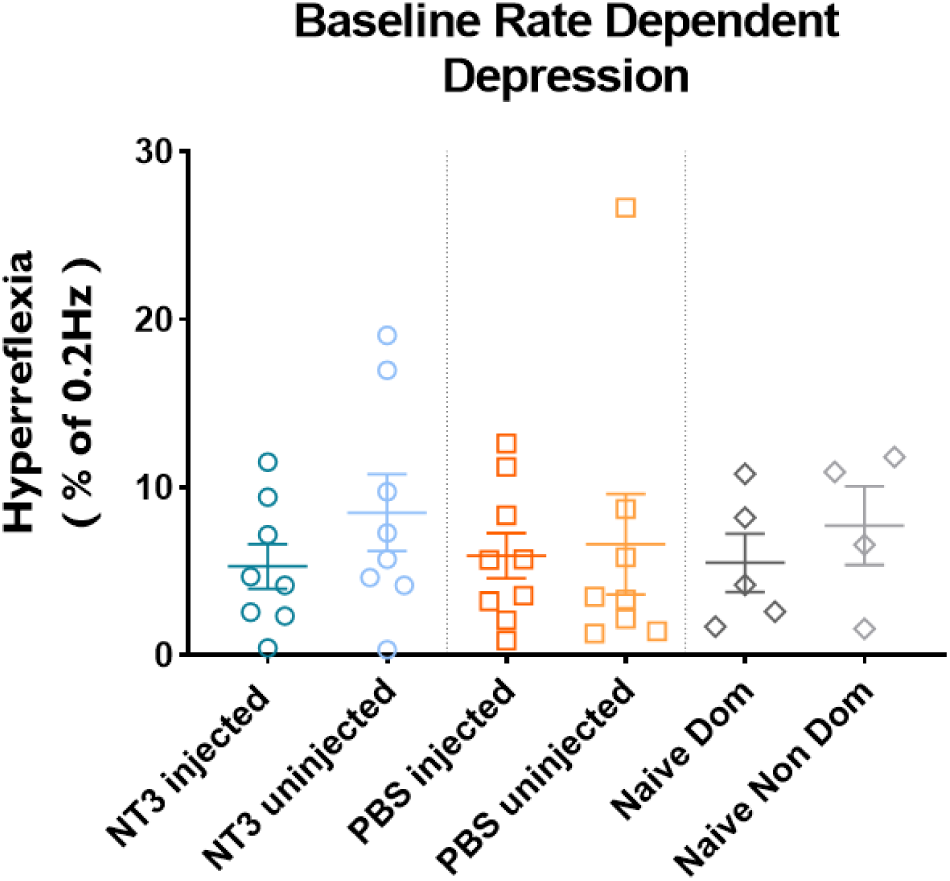
Rate dependent depression was present in all animals prior to injury. The mean amplitude of the H wave at 5 Hz stimulation ranged from 5.2 – 8.5 percent of the amplitude of the H wave during 0.2Hz stimulation. Bar one animal within the PBS noninjected group, all animals showed rate dependent depression to a degree where the H wave was depressed to 1/5 of its size. There was no significant difference between any of the groups (One- way ANOVA, F (5, 36) = 0.3585, P=0.87)

In the injected limb there was loss of rate dependent depression following injury when compared to the naïve group (Figure 5A). This difference between naïve and injured animals indicates the formation of hyperreflexia. There was no difference in hyperreflexia in the NT3 group compared to the PBS group, and once the hyperreflexia developed it remained constant. Hyperreflexia also developed in the noninjected limb of both injured groups compared to naïve animals (Figure 5B). Inclusive of all three groups there was an effect of time in the noninjected limb, indicating that hyperreflexia did increase over time. There was no interaction between group and time in either limb.

**Figure 5.**
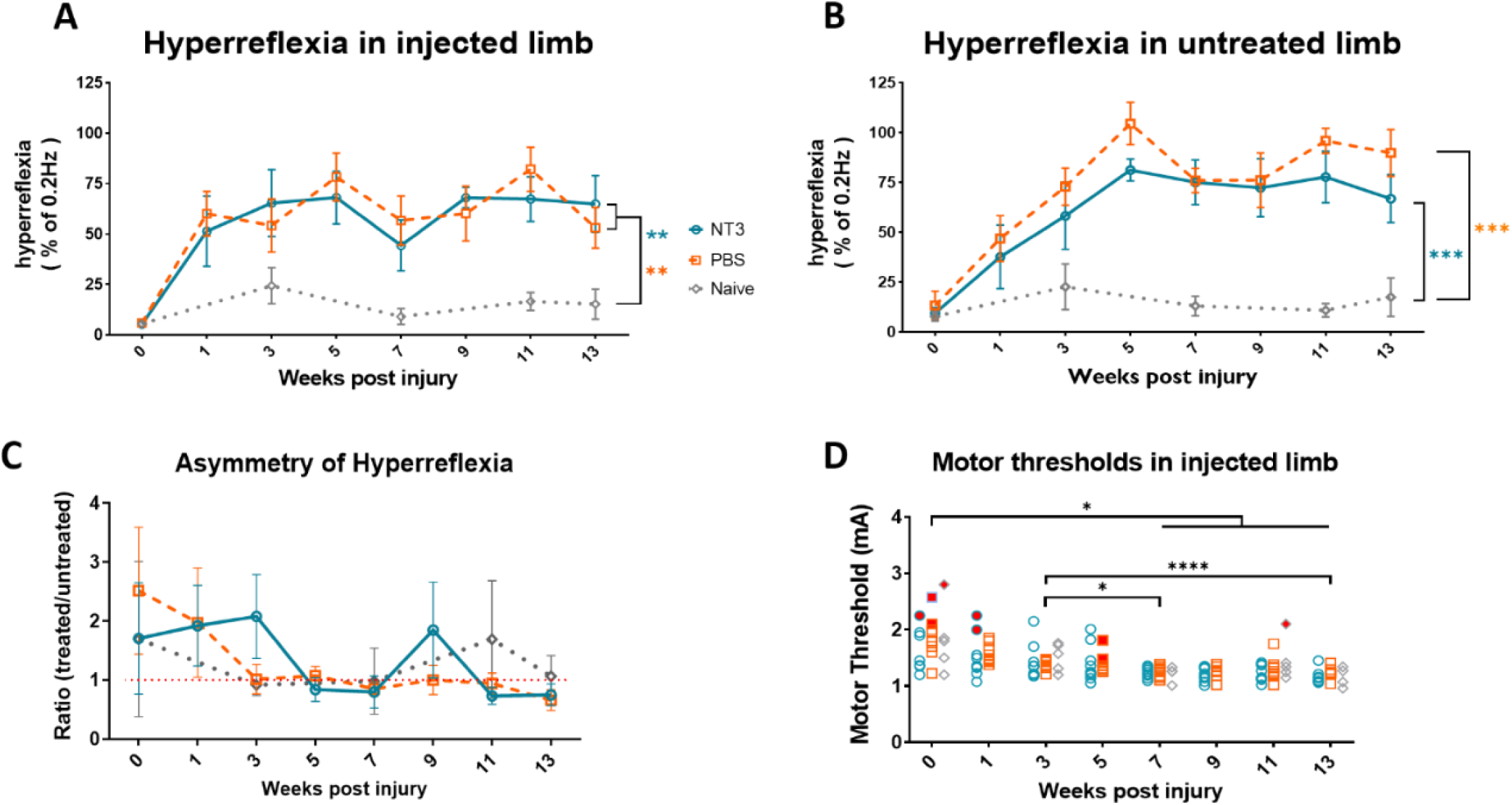
NT3 was unable to reduce hyperreflexia in distal forelimb circuitry. **A)** Contusion caused hyperreflexia in the Abductor Digiti Minimi in the injected limb (linear model, main effect of group F(2, 22.3)=6.110, p=0.008, post hoc LSD groupwise comparisons; NT3 v Naïve p=0.002 and PBS v Naïve p=0.001). Hyperreflexia in the injected limb was unaltered by NT3 treatment (effect of group, post hoc LSD for PBS vs NT3 p=0.38). There was no change over time overall (main effect of time, F(6,93.0)=1.751, p=0.118) nor an interaction between treatment and time (group x time F(10, 92.4)=0.81, p=0.62). **B)** Hyperreflexia was also present after contusion in the noninjected limb (main effect of group F(2, 18.0)=11.093, p=0.001, post hoc LSD groupwise comparisons; NT3 v Naïve p<0.0001 and PBS v Naïve p<0.0001). Hyperreflexia in the noninjected limb was unaltered by NT3 treatment (effect of group, post hoc LSD for PBS vs NT3 p=0.507). Hyperreflexia did change over time (main effect of time, F(6,85.0)=2.77, p=0.016, however post hoc LSD revealed pairwise comparison between weeks 3,7,11 &13 were p>0.15). There was again no interaction between treatment and time (group x time F(9, 85.1)=1.090, p=0.38). **C)** The ratio of hyperreflexia that developed in both forelimbs did not differ between the NT3 and PBS groups (linear model, effect of group F(2, 23.2)=0.285, p=0.76). There was no deviation away from symmetry over time overall ( linear model, main effect of time F(6, 64.7)=2.054, p=0.071) nor an interaction (linear model, group x time F(9, 62.3)=0.250, p=0.99). **D)** At baseline, the motor thresholds of the ADM muscle in injected limbs were comparable amongst all three groups; NT3=1.54 ±0.31 mA, PBS=1.64 ±0.25 mA and Naïve=1.59 ±0.30 mA (one way ANOVA of baseline values only, F(2, 14) = 0.2092, P=0.81). Over the duration of the study there was a main effect of time with decreased motor thresholds in all three groups at chronic time points compared to earlier (linear model, main effect of time F(6,68.9)=5.87, p<0.0001, LSD post hoc comparisons involving all groups; week 1 vs weeks 5-13 all p<0.05, week 3 vs week 7 p=0.012, week 3 vs week 11 p=0.063 and Week 3 vs week 13 p<0.001). NT3 treatment did not alter motor thresholds as there was no difference between groups (linear model, main effect of group F(2, 17.1)=2.140, p=0.15). Analysis performed on data with bleeding sessions, indicated by red dots, excluded. For all panels means ± S.E.M. *=P<0.05, **=p<0.01, ***=p<0.001. For NT3 and PBS groups n=7-9, for naïve n=3-5.

Since the NT3 treatment was only injected into the previously dominant forelimb this created an internal control to assess treatment effects within each animal. Overall there was no difference in the hyperreflexia in the injected over the noninjected limb of NT3 treated animals when compared to PBS treated animals, which should in theory have remained equal in both limbs (Figure 5C).

At baseline the motor threshold of the injected ADM (the stimulus required to generate a motor response) was consistent amongst all three groups (Figure 5D). The motor thresholds were comparable between both injured groups and naive animals overall, however there was a main effect of time indicating a general reduction in the motor thresholds as the study progressed. Due to the presence of bleeding from the recording site, a few individual recordings of the reflex recordings and associated motor thresholds were excluded. These are indicated by red dots in Figure 5D.

There was a shift in the latency of the H wave during 6^th^ to 20^th^ stimulus at 5Hz stimulation compared to 0.2Hz stimulation (Figure 6). By measuring the time taken for the following parameters of the H wave to occur; end of stimulus artefact, H wave onset and H wave offset, it was possible to measure both the latency of the H wave and the duration (Figure 6A). In both injured and intact animals the latency of the H wave increased during the 5Hz stimulation (Figure 6B). After injury the duration of the H wave remained comparable during 0.2Hz and 5Hz stimulation, however there was a decrease in duration in naives (Figure 6C). This was primarily because these animals had intact rate dependent depression and the H wave was barely measurable. There was no change in the offset of the H wave at either frequency (Figure 6D).

**Figure 6.**
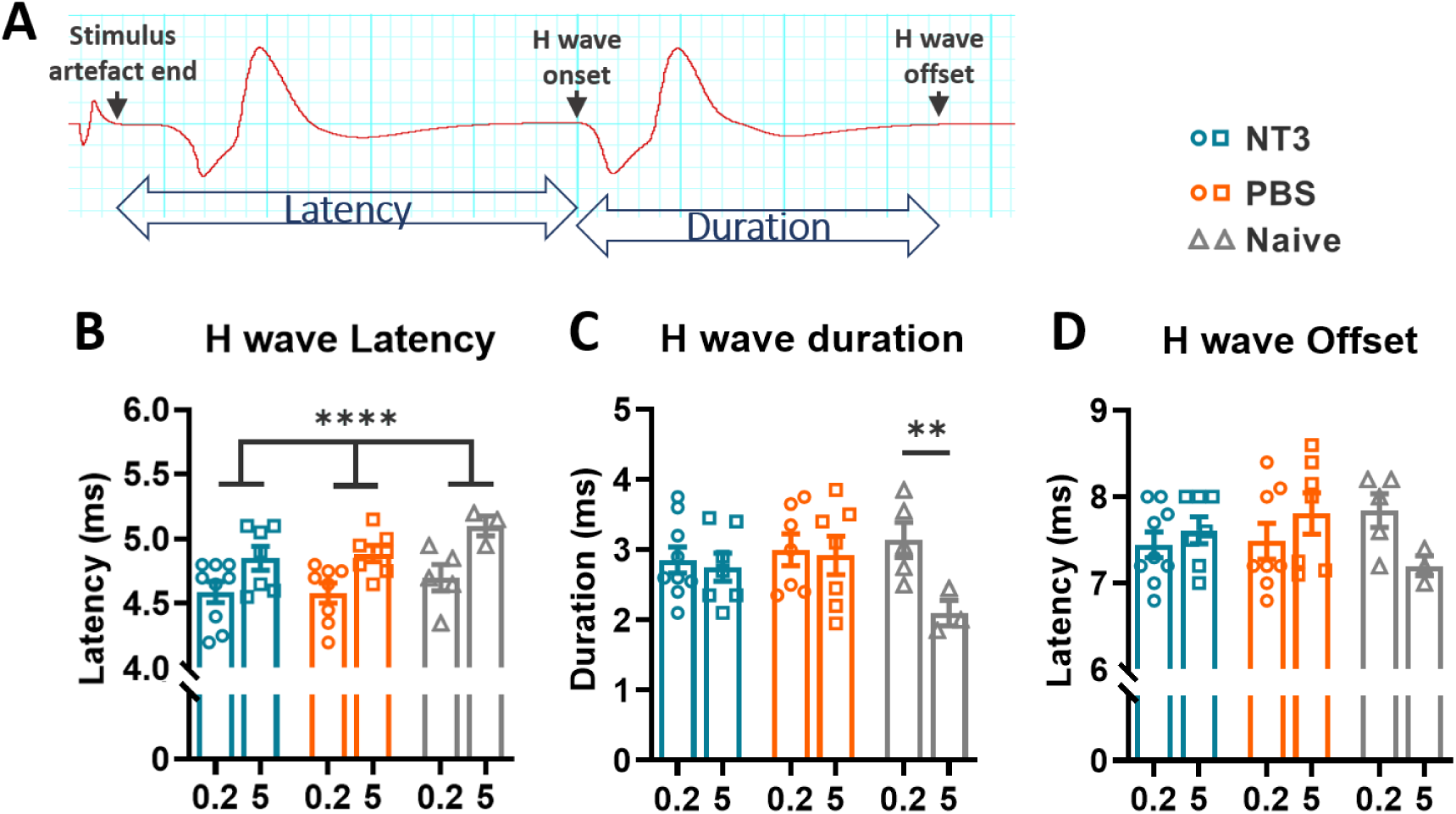
Latency of the H wave increases during 5Hz stimulation. **A)** Representative CMAP trace at 0.2Hz stimulation showing the location of measurements taken to calculate H wave latency and duration. **B)** Average latency of the H wave from the 6th to 20th stimulus increased during 5Hz compared to during 0.2Hz stimulation (mixed model effect of frequency, F(1, 14) = 97.08, p<0.0001). There was no change induced by injury or by NT3 treatment (effect of treatment, F (2, 19) = 1.070, p=0.36; interaction between frequency x treatment F (2, 14) = 0.5570, p=0.58. **C)** In Naïve animals, average duration of the H wave during the 6th to 20th stimulus decreased during 5Hz compared to 0.2Hz stimulation (mixed model, frequency x treatment, F (2, 14) = 4.94, p<0.024, effect of frequency F (1, 14) = 12.2, p=0.0036, post hoc LSD of 0.2Hz vs 5 Hz for; Naive p=0.0015, NT3 p=0.36 & PBS p=0.68). **D)** There was no change in the offset of the H wave (mixed model, effect of treatment,, F (2, 21) = 0.7450, p=0.48; effect of frequency, F (2, 21) = 0.7450, p=0.51). N for NT3=7-9, for PBS N=7-8 and sham N=5-3

### Forelimb sensorimotor function following contusion injury was slightly greater in rats treated with NT3

Initially animals were observed during locomotion in an open field each week to assess their recovery after injury, and later to identify any muscle spasms during free movement. Throughout this task and during other the other behavioural assessments below, there was no obvious muscle spasms in either forelimbs or hindlimbs nor any obvious signs of spasticity related gait disturbances as previously reported after bilateral pyramidotomy ^57^.

Locomotion across an irregular horizonal ladder tests the sensorimotor function of both forelimbs. All paw placements were scored according to a detailed seven point scale that differentiates between slip based and corrective based stepping errors ^67^. A full description of this scale is contained in the methods section but briefly; scores of two and below indicate slips, scores above three represent replacement steps and errors with paw placement, and finally a score of six indicate an error free step. Whilst animals were able to remain upright and locomote minimally around the home cage within one week of SCI, all animals were unable to make any weight supported steps on the horizontal ladder and were not formally assessed at this time point. Within two weeks animals began to traverse the ladder. Footage of this task was analysed after the ELISA results revealed that only three animals had raised NT3 serum levels and NT3 in the biceps was varied. Given this it was decided to only analyse the final time point at 12 weeks post injury, when any sustained benefit of NT3 would be apparent, with treatment effects identified by comparing to naïve animals. Blinding was maintained as during analysis only the individual animal numbers were used and these were not related back to treatment administered until all scores were collected.

There was a significant increase in the number of slip based errors 12 weeks post injury when compared to compared to naïve controls (Figure 7A). The errors occurred for both injected and noninjected forelimbs and indicates that SCI did induce a bilateral deficit in forelimb placement and locomotion across a horizontal ladder chronically. NT3 treatment did not reduce this deficit compared to the PBS group. There was also no improvement in the injected compared to the noninjected forelimb in the NT3 group. Regarding the total number of erroneous steps after injury, there was evidence that SCI caused a sustained deficit as shown by the PBS group compared to naïve (Figure 7B). There was no difference between the NT3 and PBS groups in the number of steps taken. SCI resulted in more steps by both injected and noninjected limbs required to cross the ladder (Figure 7C). A full analysis of each type of error is provided in (Figure 8). Most noticeably, NT3 treatment was able to reduce the number of minor slips compared to PBS (Figure 8C), whilst increasing the proportion of steps that needed correction during placement, a score of four (Figure 8E). This could indicate a beneficial effect of NT3 whereby improved sensorimotor connections might prevent the number of minor slips but these steps are not error free and require corrective movements in order to maintain locomotion.

**Figure 7.**
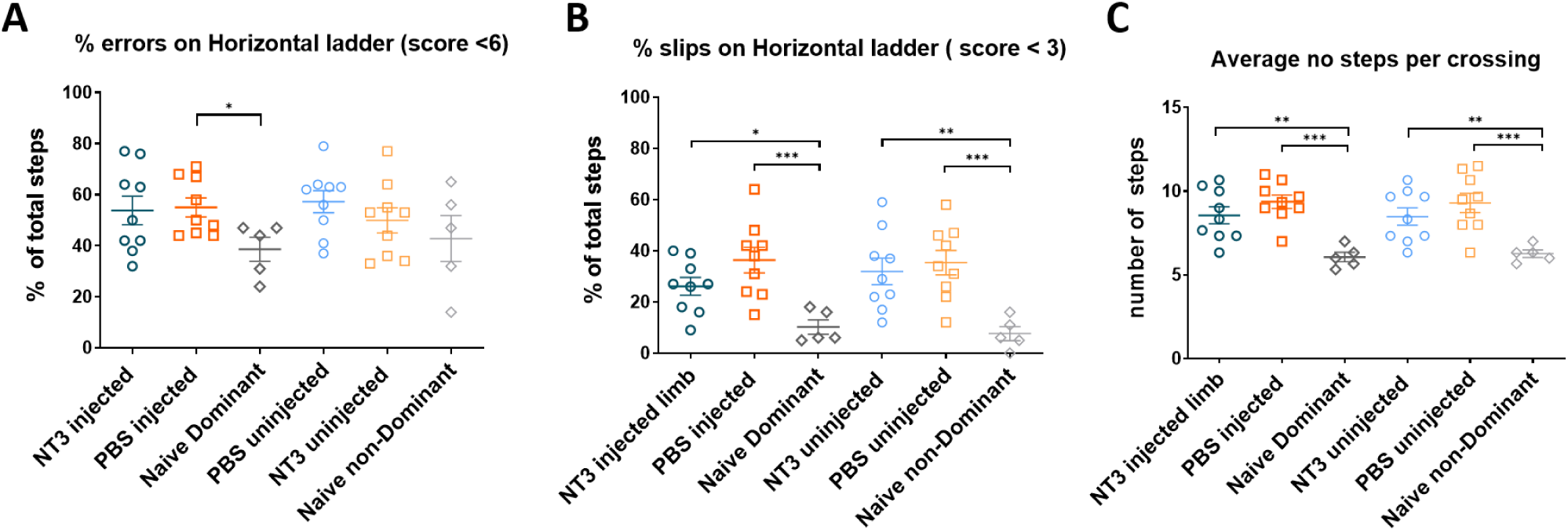
Horizontal ladder deficits were not normalised by NT3 at 12 wpi. **A)** The injury caused a significant increase in the number of slip like errors compared to naïve animals bilaterally (One way ANOVA, F(5, 40) = 6.033, P=0.0003, post hoc Fisher’s LSD for injected limbs: NT3 vs Naive p=0.031, PBS vs Naive p=0.0007, post hoc Fisher’s LSD for noninjected limbs: NT3 vs Naive p=0.0015, PBS vs Naive p=0.0004). There was no change in the number of slips by the injected limb for the NT3 group compared to PBS group (26.1 ± 10.4% and 36.3 ± 14.9% respectively, post hoc LSD p=0.0973). There was no difference between errors by the injected and noninjected limbs for either treatment groups (post hoc LSD, injected vs noninjected for; NT3 p=0.34, PBS p=0.89, naïve p=0.74) **B)** The PBS injected forelimb made slightly more placement errors, scored 0-5, in treated forelimb compared to naïve, whilst NT3 injected forelimbs made a comparable amount of errors as uninjured animals (One way ANOVA, F (5, 40) = 1.601, P<0.18, post hoc LSD for injected limbs: NT3 vs Naive p=0.068, PBS vs Naive p= 0.048 post hoc LSD for noninjected limbs NT3 vs Naive p=0.38, PBS vs Naive p= 0.14). NT3 was unable to reduce the number of erroneous steps compared to PBS for either limb (injected limb NT3 vs PBS p=0.86 and noninjected limb NT3 vs PBS p=0.39). There was no difference between injected and noninjected limbs for either treatment groups (post hoc LSD test injected vs noninjected for; NT3 p=0.57 and PBS p=0.75, respectively). **C)** Injured animals took more steps to cross the ladder compared to naïve animals (one way ANOVA, F (5, 40) = 6.967, P<0.0001, post hoc LSD for injected limb; NT3 vs naive p=0.0024, PBS vs naive p=0.0001 and for noninjected limb NT3 vs naive p=0.0062, PBS vs naive p=0.0003). There was no difference between the NT3 and PBS groups in either limb (NT3 injected vs PBS injected p=0.21, NT3 noninjected vs PBS noninjected p=0.22). For NT3 and PBS groups n=9, for naïve n=5. Error bars are S.E.M, scores averaged from a total of 3 runs. Baseline errors during horizontal ladder were not included as a covariate. * indicate significance based on main effect; * indicates P<0.05, ** indicate P<0.01, *** indicate P<0.001.

**Figure 8.**
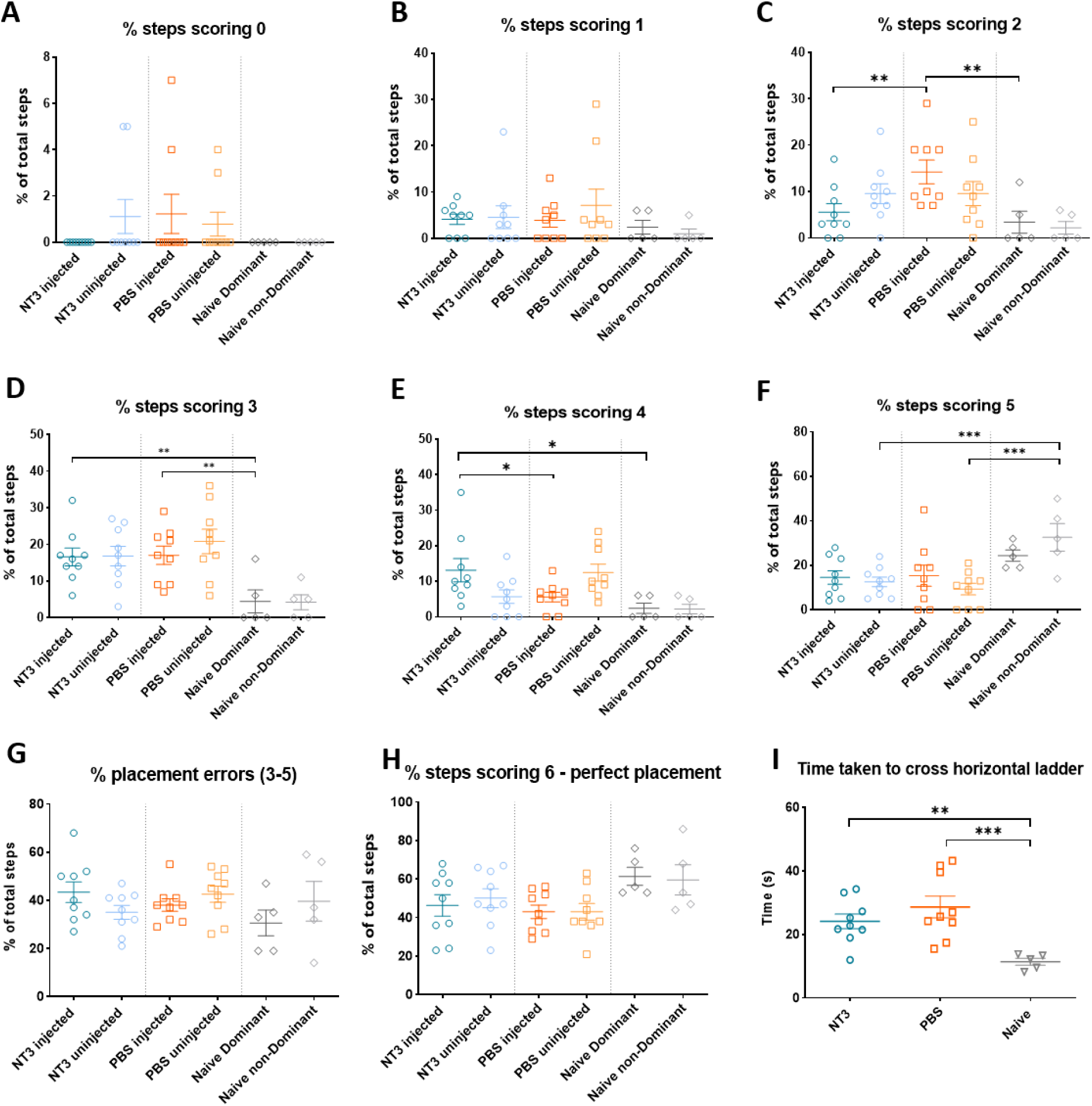
Comprehensive scoring of Horizontal ladder forepaw errors. **A)** There was no difference in the number of total missed steps which stopped locomotion (one way ANOVA F(5,40)=0.94, p=0.46). **B)** There was no difference in the number of deep slips up to the elbow (one way ANOVA F(5,40)=0.7188, p=0.61). **C)** NT3 animals performed fewer slight slips compared to PBS animals, to a level comparable with naïves (one way ANOVA F(5, 40) = 3.450, p=0.011, post hoc LSD for injected limb NT3 vs PBS p=0.0069, NT3 vs Naïve p=0.55). PBS treated animals performed more slips compared to uninjured animals (post hoc LSD for injected limb PBS vs Naïve p=0.0045). There was no difference between limbs in injured or naives (injected vs noninjected for NT3 p=0.1958, for PBS p=0.13 and naïve p=0.77) **D)** Animals performed more replacement steps following injury but this was not prevented with NT3 (one way ANOVA F(5,40)=4.923, p=0.0013, post hoc LSD for injected limb NT3 vs Naïve p=0.0086, PBS vs Naïve p=0.0066, NT3 vs PBS p=0.91; for noninjected limb NT3 vs Naïve p=0.0067, PBS vs Naïve p<0.0003, NT3 vs PBS p=0.29). **E)** The number of corrective steps were increased in NT3 but not PBS group after injury (one way ANOVA F(5,40)=4.067, p=0.0044, post hoc LSD for injected limb NT3 vs Naïve p=0.0047, PBS vs Naïve p=0.38). There was a significant increase in these steps following NT3 treatment (injected NT3 vs Injected PBS p=0.017), with more occurring in the injected limb (injected NT3 vs noninjected NT3 p=0.018). There was also significant difference between forelimbs in the PBS group (injected PBS vs noninjected PBS p=0.028) **F)** Partial placement steps were similar between the injected limbs of injured animals compared to naives, but were reduced in the uninjured limbs (one way ANOVA F(5, 40) = 4.521 p=0.0023, post hoc LSD for injected/dominant limb; NT3 vs naïve p=0.0849, PBS vs naïve p=0.11, NT3 vs PBS p=0.87, comparison between noninjected limbs; NT3 vs naïve p=0.0009, PBS vs naïve p=0.0002, NT3 vs PBS p=0.50). **G)** The percentage of perfectly placed steps was comparable amongst all groups and limbs (one way ANOVA F(5, 40) = 2.087 p=0.087). **H)** Overall there was no different in errors of placement between all three groups (one way ANOVA, F(5, 40) = 1.223 p=0.32). **I)** Animals took a longer time to cross the ladder after SCI, which was not improved with NT3 treatment ( one way ANOVA, F(2, 20) = 7.648 p=0.0034; NT3 vs naïve p= 0.0097, PBS vs naïve p=0.0009, NT3 vs. PBS p=0.2397). For NT3 and PBS groups n=9, for naïve n=5. Error bars are S.E.M, scores averaged from a total of 3 runs. Baseline errors during horizontal ladder were not included as a covariate. * indicates P<0.05, ** indicate P<0.01, *** indicate P<0.001.

### Single Pellet Reaching

Success at the single pellet reaching task requires fine sensorimotor function and proprioception to carry out the reaching, grasping and finally retrieval of small sugar pellets. In order to prevent the compensatory movement of dragging of pellets instead of grasping with the impaired forepaw, a custom device was made with a gap which separated the ledge where the pellet is presented from the reaching window (Figure 9A). At baseline all animals had achieved a similar ability to grasp pellets with one or more attempts (Figure 9B). Animals below 55 percent success rate at baseline, regardless of the number of attempts, were not included in the study. Whilst most injured animals were able to reach for pellets, three animals (n=1 for NT3 and n=2 for PBS) were excluded from the post injury analysis due to being unable to make any on target reach attempts, i.e., making contact with the pellet, at any timepoint after injury.

**Figure 9.**
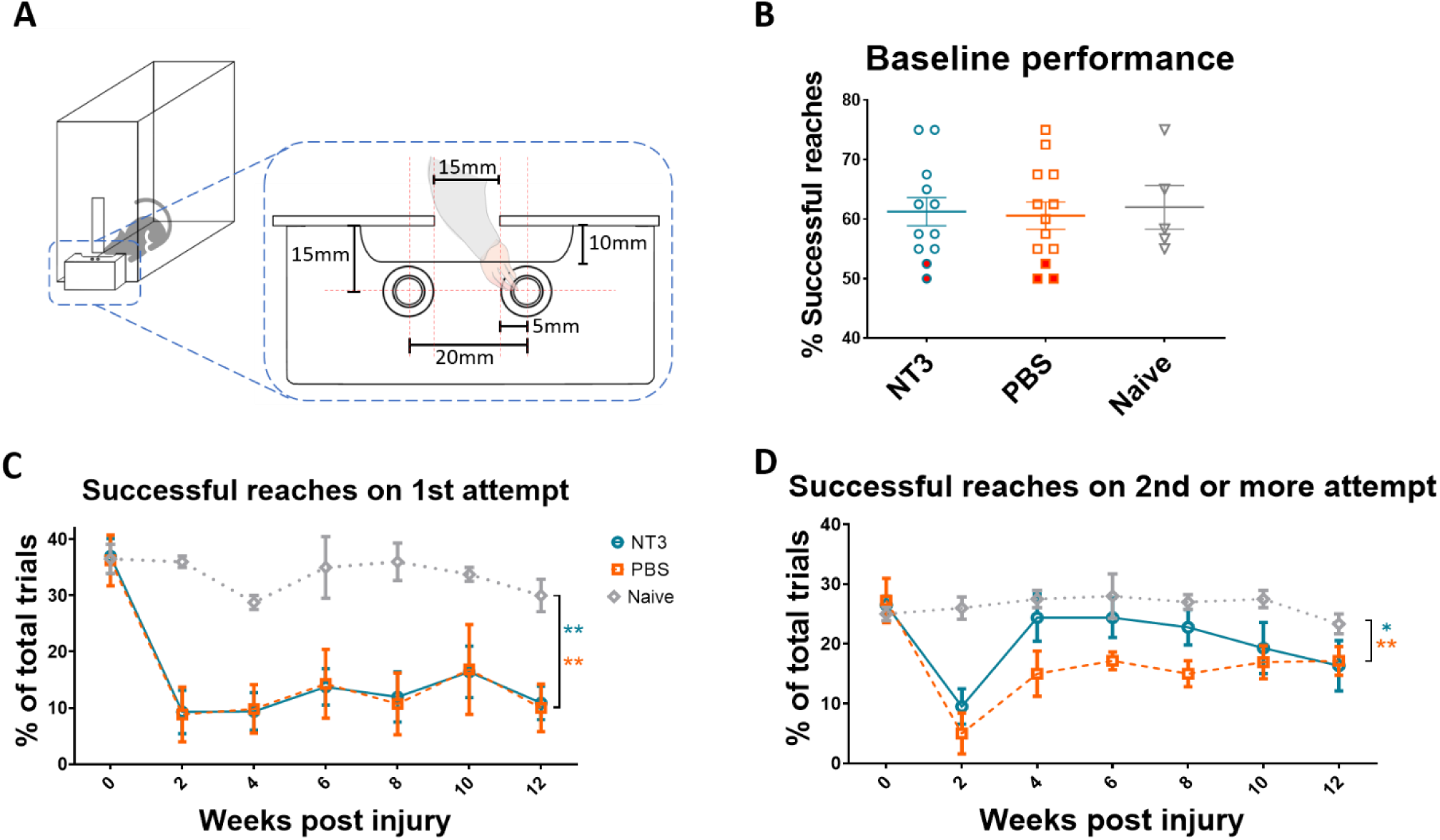
NT3 did not normalise the deficit in reach and grasp which occurs after SCI. **A)** Schematic of the pellet reaching apparatus used for the task. **B)** All three groups had similar pellet reaching performance at baseline (one-way ANOVA, F (2, 17) = 0.071, p=0.79). Red dots indicate animals that failed to reach 55% overall successful grasp rate following training and were excluded from the study prior to surgery and were excluded from the entire study. **C)** A deficit in the percentage of successes on first attempt was present after injury (effect of group, linear model, F(2, 15.6) = 8.900, p=0.003, post hoc LSD comparisons for NT3 vs Naïve p=0.0016, PBS vs Naïve p=0.0017, NT3 vs PBS p=0.94). There was no change in ability over time in any group (effect of time, linear model, F(5,16.6) = 1.2, p=0.34), nor any effect of NT3 treatment (time * group F(10, 16.79) = 0.200, p=0.074) . **D)** The deficit persisted in the percentage of reaches that were successful but required more than one attempt (effect of group, linear model, F (2, 17) 6.45, p=0.0083,NT3 vs Naïve p=0.045, PBS vs Naïve p=0.0023, NT3 vs PBS p=0.12). There was a change in the ability to complete the task with multiple attempts over time (effect of time, linear model, F(5,16.8)=7.6, p=0.0068, post hoc LSD comparisons including all groups; 2 wpi vs 4 wpi p=0.00075, 4wpi vs later time points were all p>0.43). NT3 had no significant effect on improving successful reaching after multiple attempts (effect of group*time, linear model, F (10, 17) = 1.81, p=0.13). For NT3 n=8, for PBS n=7 and for naïve n=5. Averages are from 20 trials in a single session per week. * indicate significance based on main effect; * indicates P<0.05, ** indicate P<0.01, *** indicate P<0.001.

**Figure 10.**
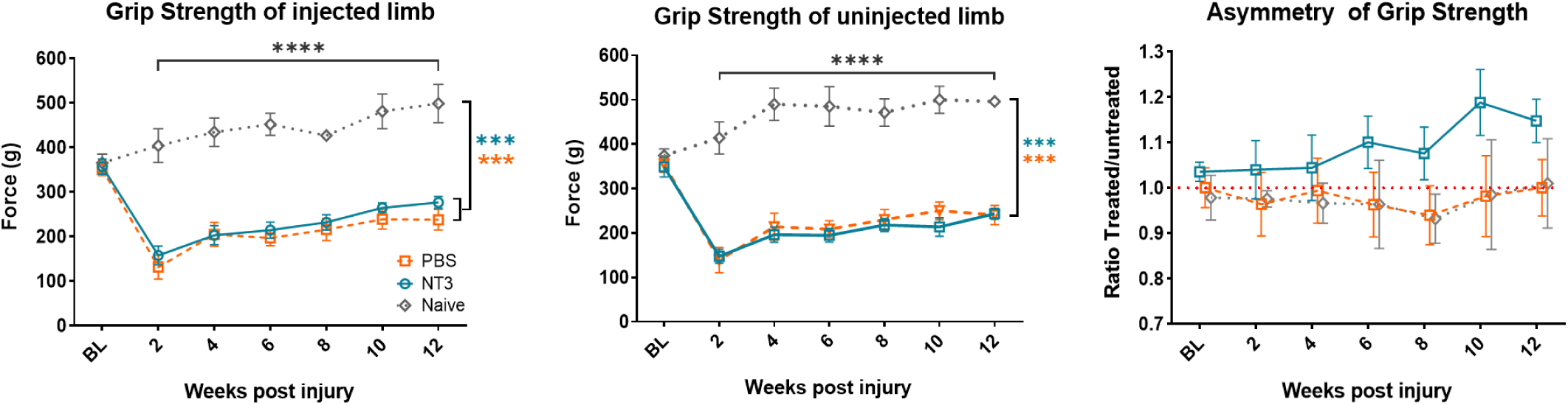
Contusion caused sustained deficits in grip strength which was not normalised by NT3 treatment. **A)** There was a comparable reduction in grip strength in the injected limb of both NT3 and PBS groups following injury, however NT3 failed to have any effect on grip strength (Linear model main effect of group,F2,18=44.0, p<0.0001, post hoc LSD comparison, NT3 vs Naïve p<0.0001, PBS vs Naïve p<0.0001, NT3 vs PBS p=0.15). Additionally there was increased grip strength in all groups as the study progressed (effect of time, linear model, F (5, 91)=12.0, p<0.001, post hoc LSD comparisons; 2 wpi vs 4 wpi p=0.002, 2wpi vs 12 wpi p<0.0001). There was no interaction between group and time post injury (effect of time x group, linear model, F (10, 91)=0.57, p=0.83). **B)** There was a similar reduction in grip strength after injury in the noninjected limb (Linear model main effect of group, F (2, 22) = 65.37, p<0.0001, post hoc LSD comparison, NT3 vs Naïve p<0.0001, PBS vs Naïve p<0.0001, NT3 vs PBS p=0.76). An effect of time was also present ( linear model, F(5, 95) = 10.30, p<0.0001, post hoc LSD comparisons; 2 wpi vs 4 wpi p=0.001, 2wpi vs 12 wpi p<0.0001). There was no interaction between group and time post injury (effect of time x group, linear model, F (10, 95) = 0.3551, p=0.96). **C)** Considering all time points together, grip strength remained symmetrical overall (main effect of group, linear model, F(2,21)= 0.007, p=0.99). However, there was weak evidence for greater grip strength in the treated limb relative to the untreated limb over time in the NT3 group relative to the other groups (group x time interaction, linear model, F(10,110)=2.049 p=0.035); however the apparent divergence at 10 and 12 weeks post injury in NT3 group was not significantly different to PBS (post hoc LSD at week 10, NT3 vs PBS p= 0.16; at 12 weeks NT3 vs PBS p=0.97. There was no overall consistent effect of time (linear mixed model, F(5,110)= 0.123, p=0.352). For NT3 n=9, for PBS n=9 and for naïve n=5. Averages are from 3 non-consecutive trials in a single session each week. * indicates P<0.05, ** indicate P<0.01, *** indicate P<0.001

**Figure 11.**
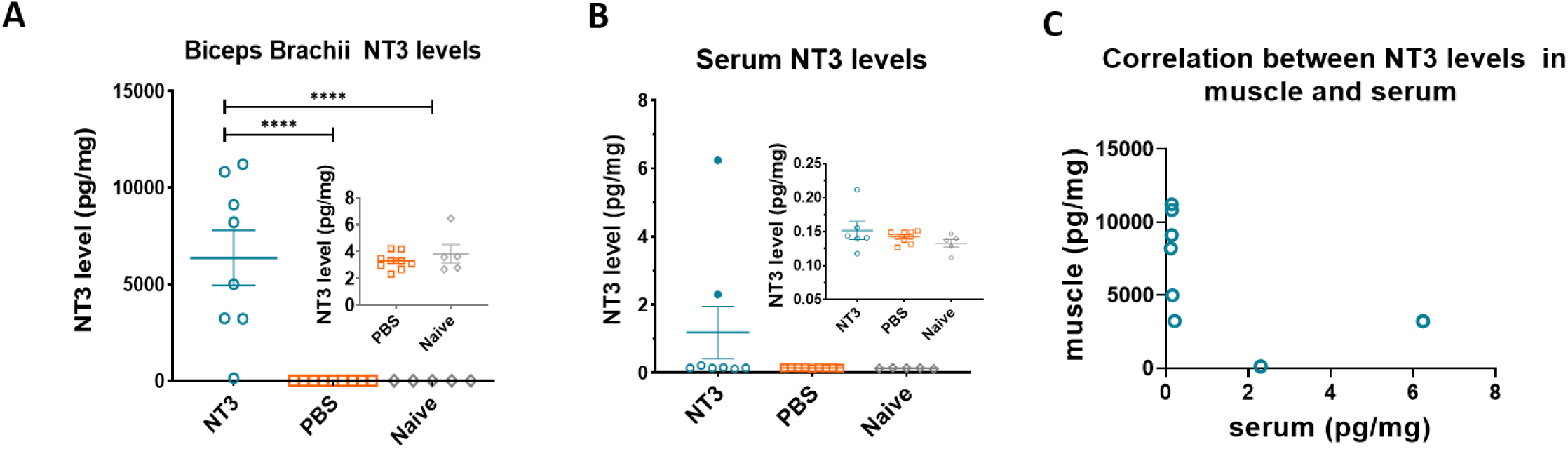
Intramuscular injections of AAV-NT3 raised NT3 protein in muscles but not consistently in serum. A) Intramuscular injection of AAV-NT3 into the biceps brachii resulted in a significant increase in the amount of NT3 protein produced, 6366 ±4050 pg/mg, compared to both PBS and Naïve, 3.28 ± 0.63pg/mg and 3.82 ±1.54 pg/mg respectively (one way ANOVA, main effect of group F(2, 19) = 17.06, p<0.0001, post hoc LSD NT3 vs PBS p <0.0001, NT3 vs Naïve p=0.0002). SCI followed by intramuscular injections of PBS did not alter the level of NT3 in biceps brachii, shown in insert (post hoc LSD, PBS vs Naïve p=0.99). One animal in the NT3 treated group had a bicep level of 128.6 pg/ml NT3. B) Serum levels in the treated animals were inconsistent and as a cohort were not raised above PBS or Naïve groups (F (2, 19) = 1.583, p=0.23). Contusion injury alone did not alter serum NT3 levels chronically, 0.14 ± 0.01pg/mg and 0.13 pg/mg ± 0.01 pg/ml in PBS and Naïve respectively (see insert, post hoc LSD PBS vs Naive p=0.99). Only three animals had raised serum levels of 0.21, 2.30 & 6.23 pg/ml. C) Surprisingly there was no correlation between the level of NT3 in the serum and Biceps Brachii (r=-0.5371, p=0.11)

Spinal cord injury alone caused a significant deficit in number of pellets reached on the first attempt compared to naives (Figure 9 C). This deficit was not improved with NT3, and the percentage of hits remained comparable between the NT3 and PBS groups. Overall there was no change in pellet reaching over time nor any interaction between treatment and time post injury.

There was a similar deficit after injury in the number of pellets that were successfully grasped and retrieved following more than one attempt. Again, the NT3 and PBS groups did not differ overall and both had significantly reduced performance compared to naïves. A main effect of time was present, indicating that on average animals retrieved a higher number of pellets requiring multiple reach attempts as the study progressed. Taking both injured groups and naives into consideration, there was a significant increase in performance between two weeks and four weeks, however performance at subsequent time points remained comparable to week 4. As before, there was no interaction between the group and time post injury and as a result, no treatment effect for NT3.

### Grip strength

Bilateral cervical contusion caused a bilateral loss in grip strength. Regarding the injected forelimbs (Figure 10A), NT3 treatment did not have any effect in reducing the grip strength deficit in the injected forelimb compared to PBS group. Both injured groups as well as naïve animals increased their grip strength over the duration of the study; possible explanations for this are explored in the discussion. The ratio of grip strength in the injected forelimb compared to noninjected of grip strength was assessed (Figure 10B). During the twelve weeks following injury the ratio of grip strength in the injected and noninjected limbs of both groups remained comparable to that of naives. Overall, there was no change in the ratio of grip strength in injected compared to noninjected over time, and whilst the PBS and NT3 groups at late time points appear to diverge, there was only weak evidence for this change. These results indicate that NT3 treatment had no effect of altering grip strength in the injected limb in relation to the noninjected limb, nor is able to rescue the significant drop in grip strength after SCI.

### Intramuscular injection of AAV1-CMV-NT3 increased NT3 levels in Muscle but not consistently in serum

Animals were perfused at 14 weeks after injury and the medial head of the biceps brachii of the injected forelimb, as well as blood, were taken to assess level of NT3 protein in these tissues. NT3 levels were expressed as a proportion of total protein present in the tissue to account for differences in protein extraction.

Intramuscular injections into the biceps brachii and other musculature resulted in an increase in NT3 protein within the muscle by 14 wpi (Figure 11A). On average AAV-NT3 increased NT3 protein levels approximately 1600 fold from levels seen in PBS treated animals and naives. SCI plus PBS injection did not alter the amount of NT3, which was comparable to levels in naives. One animal which was injected with AAV-NT3 had a much lower NT3 level of 128.6 pg/mg in the medial head, which was 50 fold lower than the average for the group.

In general, AAV-NT3 did not consistently raise serum levels throughout the cohort (Figure 11B). Only three animals had raised NT3 levels which corresponded to a fold increase of 1.5x, 16.5x and 44x above the average level in uninjured animals. The rest of the AAV-NT3 treated animals had levels in a similar range of untreated animals and therefore AAV-NT3 was unable to generate a significant increase in serum levels across the whole cohort.

Unexpectedly, overall the levels in serum did not correlate with levels in the bicep muscle (Figure 11C). The two animals which had the lowest amounts of NT3 in the biceps brachii muscle had the highest elevation in serum levels. It should be noted that only a single head of the biceps brachii was analysed, and many additional muscles were also injected which may explain this difference. Explanations as to why serum levels were not raised in all animals are explored in the Discussion.

Ex vivo MRI reveals disruption of white matter tracts and cavity formation.

T2 weighted MRI images were generated of the cervical cord in order to compare lesion size amongst the two groups (Figure 12). On T2 weighted images proton rich liquids, like cerebrospinal fluid, are represented by a hyperintense signal and appear bright white. Grey matter appears as a pale grey signal and white matter appears a darker grey. With an isotropic voxel size of 50μm x 50μm x 50μm, the characteristic morphology of the grey matter at the different cervical levels are clearly defined (Figure 12B & D). The 225 kDyn contusion injury generated a substantial lesion resulting in both the formation of a cavity, shown as a black hypointense signal, as well as abnormal tissue surrounding the cavity (Figure 12C & E-G). This abnormal tissue appears as hyperintense signal, bright white on the T2 weighted image, and can be distinguished from the spared white matter laterally. At 14 weeks after injury spinal cord volume was significantly reduced throughout the lesion site (Figure 13A). This atrophy of the spinal cord extended throughout the lesion site and beyond the area of cavitation. Owing to the presence of hypo and hyperintense throughout the lesion it was not possible to accurately measure the size of the lesion directly using the software available. Instead the amount of presumptive spared white matter, defined as white matter with a similar signal intensity to that of the white matter in the lateral funiculus region at the rostral and caudal most transverse section, was measured. SCI caused a reduction in presumptive spared white matter when compared to naïve (Figure 13B). There was approximately a 3.5-fold reduction in the total amount of presumptive white matter present around the epicentre, 0.82 ±0.12 mm^2^ in PBS group and 0.92 ±0.09 mm^2^, compared to 3.01±0.18 mm^2^ in naives. Loss of white matter extended for 5mm rostral and caudal to the epicentre. Interestingly there was a small but significant difference in the amount of putative white matter in the NT3 treated animals surrounding the epicentre and up to 1mm caudally.

**Figure 12.**
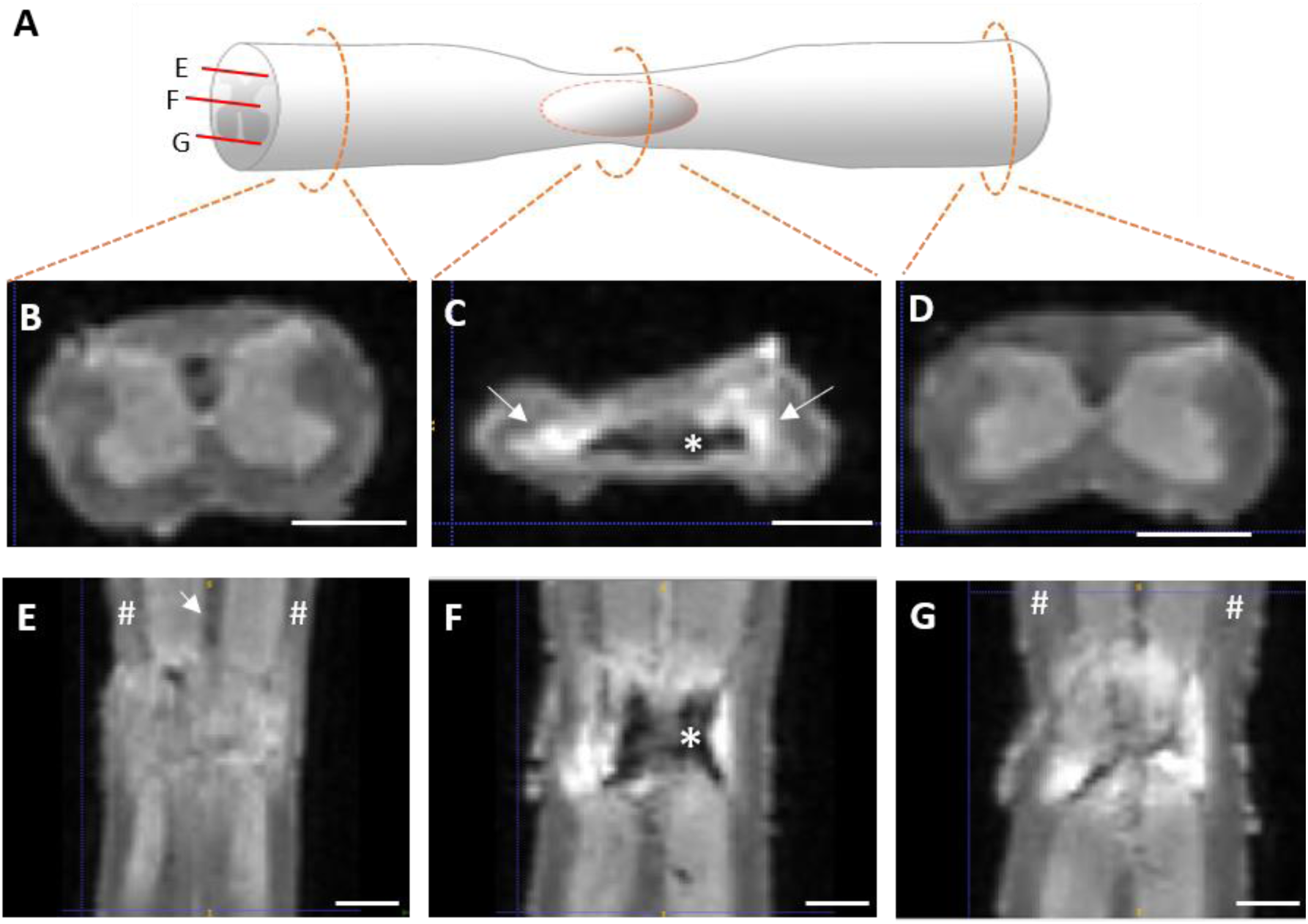
Ex-vivo T2 MRI reveals cavitation of spinal cord and disruption of white matter tracts. **A)**Schematic of injured spinal cord showing approximate locations and orientations of subsequent MRI views. Rostral is left, caudal is right. **B-D)** Cross sections through the cord at C6, C7 incorporating the epicentre and C8 spinal levels respectively. Cavitation within the epicentre (*) is apparent as a region of hypointense darker signal. This is surrounded by regions of hyperintense signal (long arrow), which are more apparent in the lateral borders of the lesion. **E-G)** Horizontal sections of the lesion at the dorsal, intermediate, and ventral levels. The lesion caused disruption to the dorsal column (short arrow) and the lateral funiculus (#) which appears more severe in the dorsal compared to the ventral region (compare # in E with G). Scale bar =1mm throughout.

Extent of cavitation was assessed using the continuous hypointense signal, present within the centre of the lesioned spinal cord, as a marker. There was progressive increase in the cross sectional area of the cavity towards the epicentre, which was defined as the single spinal cord level with the highest area of cavitation (Figure 13C). Taking the whole lesion into account there was no difference in the volume of cavitation between groups. Total cavity volume was comparable between groups. When looking at the extent of the lesion in either the rostral or caudal direction from the epicentre, cavitation extended less rostrally in NT3 treated animals compared to PBS treated animals (Figure 13D). The extent of the cavity caudally to the injury was comparable (Figure 13D).

Both groups displayed lesions of comparable size (Figure 13C). This corroborates the fact that there was no difference between displacement of spinal cord or the force of the impact between the two groups (Figure 14A & B). In both injured groups there was no strong evidence of a correlation between the displacement of the spinal cord during contusion and spared white matter at the epicentre (Figure 13 C) (*n.b.*, variation in displacement was very small). The epicentre from each animal and the automatically generated mask used to calculate presumed spared white matter is presented in Figure 15.

**Figure 13.**
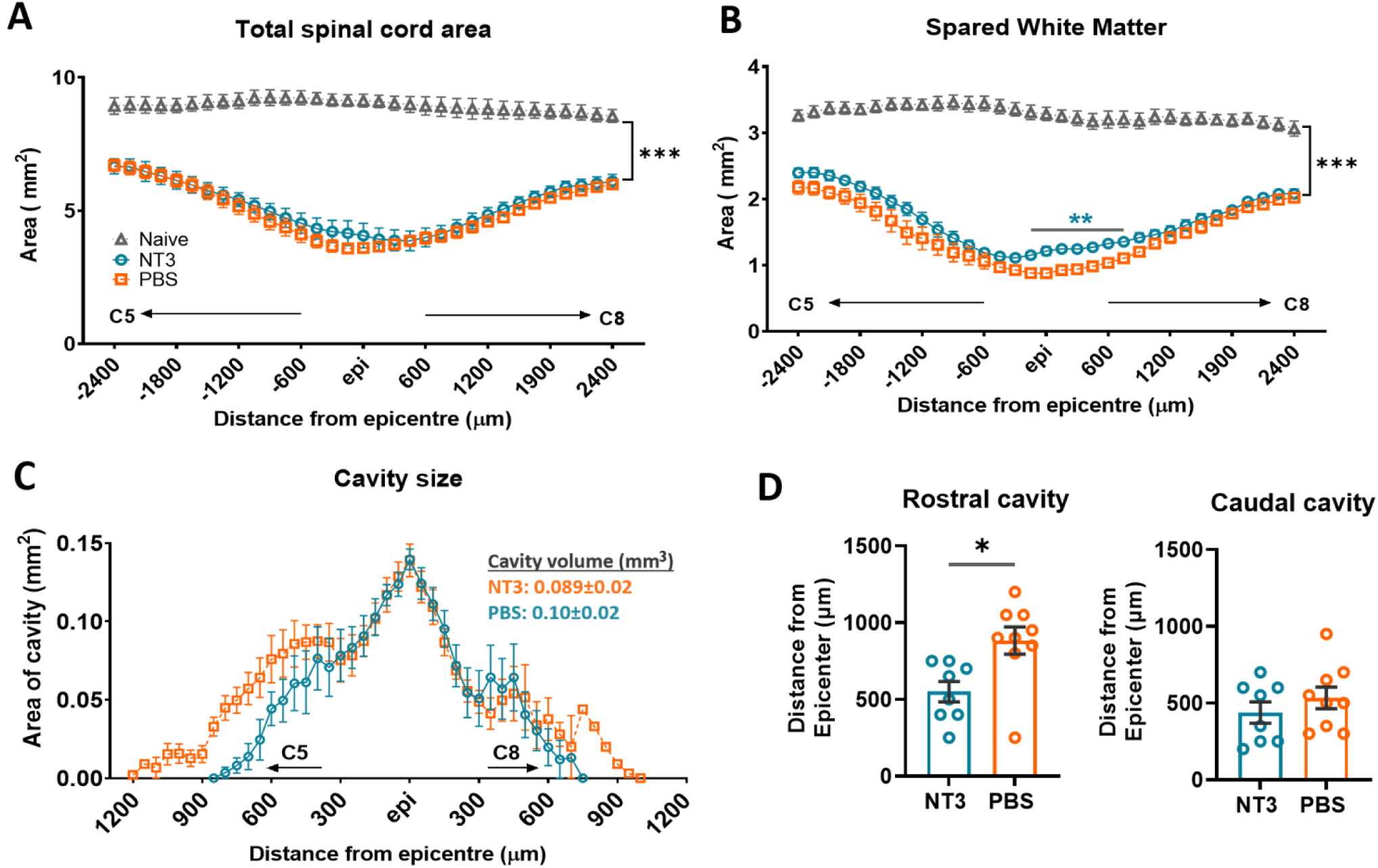
Ex-vivo MRI reveals compaction of tissue around lesion site and extensive white matter loss. **A)** Contusion caused a reduction in total spinal cord area, inclusive of cavitation, throughout the observable lesion in both injured groups (effect of group, Two way RM ANOVA, F(2, 19) = 66.00, p<0.0001). There was an interaction between group and the spinal level ( F(64, 608) = 14.00, p= <0.0001). Spinal cord area was reduced up to 2.4mm rostral and caudal from the epicentre (effect of group * level, rostral most region; NT3 vs naïve p=0.0004, PBS vs naïve p=0.0004, at epicentre ; NT3 vs naïve p<0.0001, PBS vs naïve p<0.0001 and caudal most region NT3 vs naïve p<0.0001, PBS vs naïve p<0.0001). Total cord area throughout the lesion was similar between NT3 and PBS groups (effect of group excluding naïve, Two way RM ANOVA F(1, 15) = 0.40, p=0.56). **B)** Spared white matter was significantly decreased throughout the lesion in both injured groups compared to naïves (effect of group, Two way RM ANOVA, F(2, 19) = 207.5, P<0.0001). There was an interaction between group and spinal level (F (64, 608) = 10.20, p<0.0001, post hoc LSD rostral most region NT3 vs naïve p<0.0001, PBS vs naïve p<0.0001 ; at epicentre, NT3 vs naïve p<0.0001, PBS vs naïve p<0.0001; and caudal most region NT3 vs naïve p=0.0002, PBS vs naïve p<0.0001). There was significantly increased spared white matter around the epicentre site in the NT3 group compared to PBS (effect of group * level, post hoc LSD, NT3 vs PBS at epicentre p=0.0016, at 900um caudal p=0.045, comparison for region in between were all p<0.005). **C)** Cavity size was estimated by measuring the continuous hypointense signal within the centre of the lesion. Over the entire lesion, there was no difference in the distribution of cavitation ( Two way RM ANOVA,effect of group, F (1, 15) = 3.245, p<0.091). Total cavity volume was comparable between groups (Unpaired Student T-test, p=0.10). **D)** The maximal rostral extent of cavitation from the epicentre was greater in PBS than NT3 rats (Unpaired Student T-test, P=0.01). There was comparable extent of the cavity in the caudal direction between the two injured groups (Unpaired Student T-test, P=0.35). No covariates were included in these analyses.

**Figure 14.**
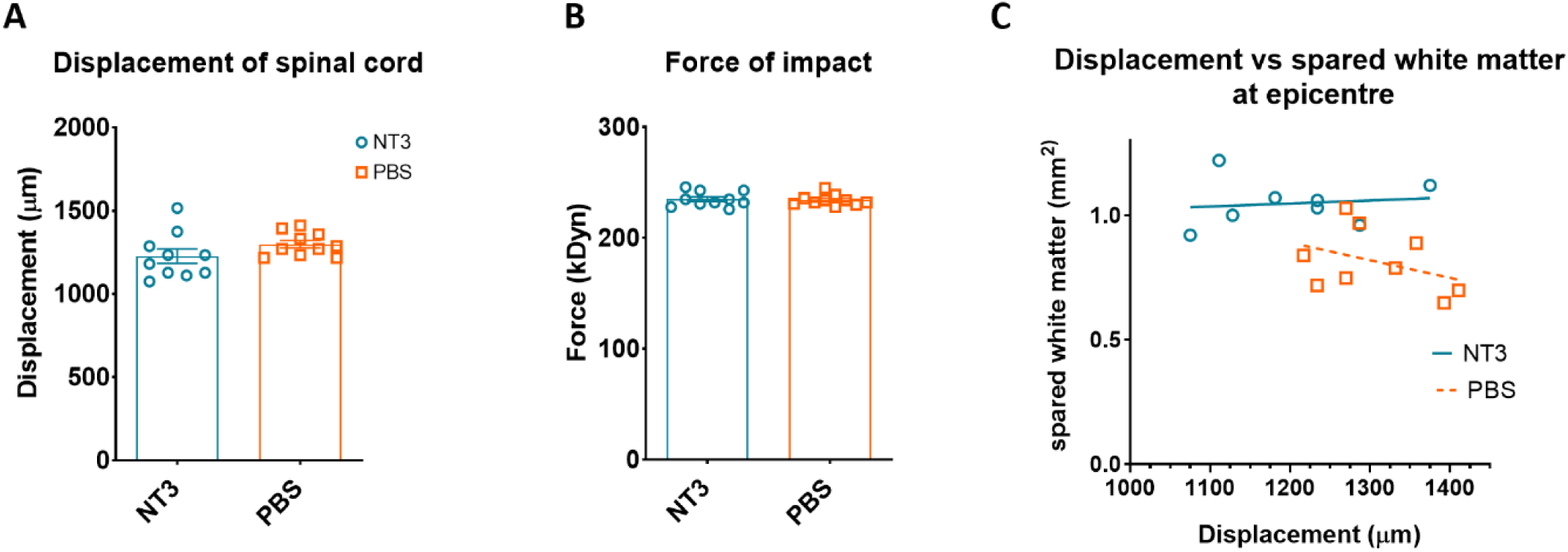
Both treatment groups received equal C5-C6 moderate contusion. **A)** The resultant displacement of the spinal cord **B)** and the force of impact, as measured by the IH infinite Horizon Device, showed no difference between the two groups. Average displacement in NT3 (1227±136 μm, n=10) and PBS (1299±71 μm, n=10) showed no difference (Unpaired Student T-test, P=0.17). Average force in NT3 (235.1±6.7 kDyn, n=10) and PBS (234.1±4.9 kDyn, n=10) showed no difference (Unpaired Student T-test, P=0.71). **C)** There was no correlation between spared white matter within the lesion epicentre and injury displacement in NT3 group (r=0.129, p=0.76, R^2^=0.016) or the PBS group (r=-0.382, p=0.31, R^2^=0.146). Error bars are S.E.M, n=8 and 9 for NT3 and PBS groups respectively, n=5 for naïve.

**Figure 15.**
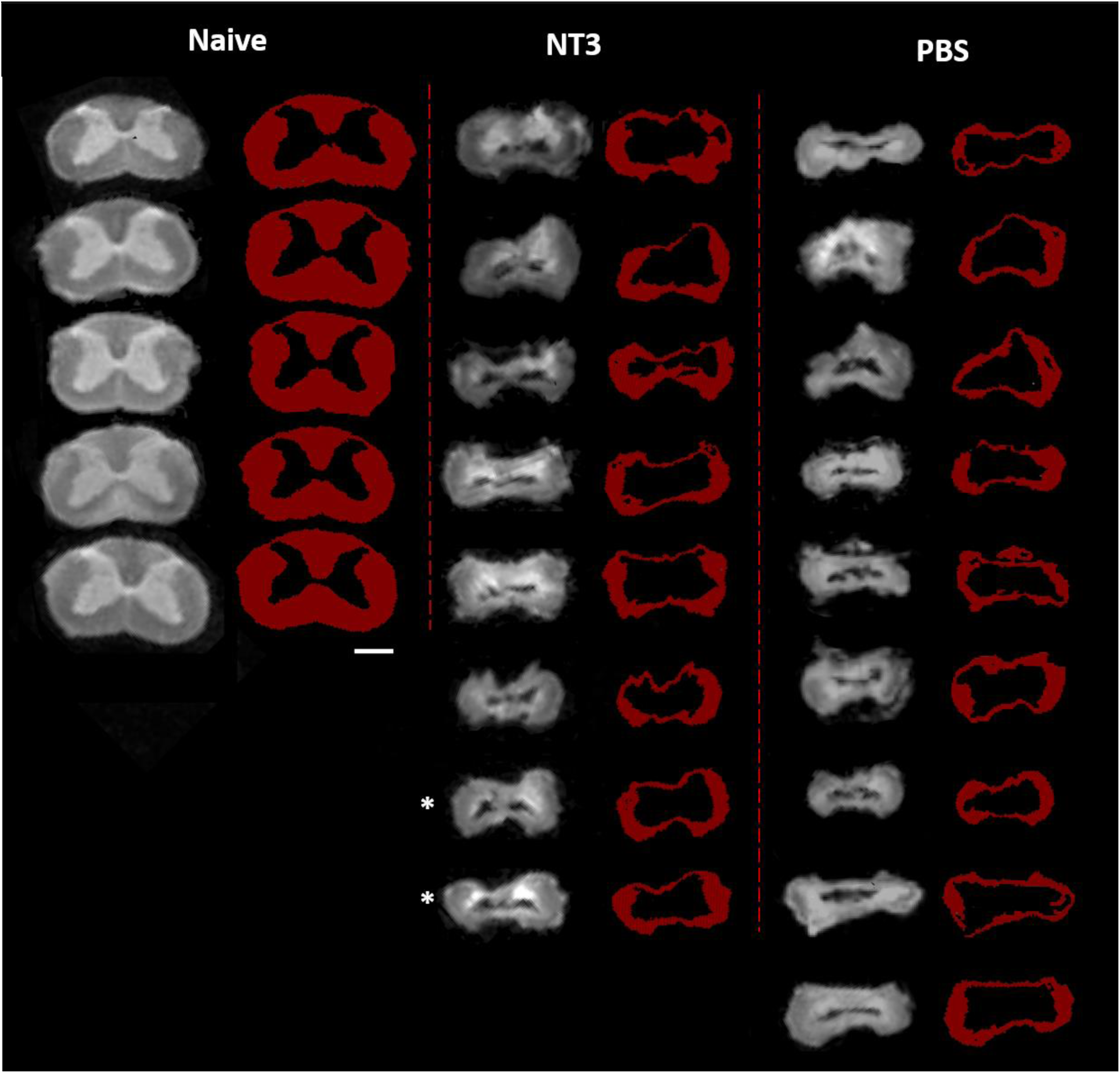
T2 weighted MRI and threshold images used during analysis of presumptive spared white matter. Greyscale images show T2 weighted MRI scans at the epicentre of all injured animals and the corresponding spinal cord level in naïve animals. Red images show the automatically segmented image which was used to calculate the area of presumptive spared white matter. There is noticeable compaction of the spinal cord after injury compared to naïve. The two animals with markedly raised NT3 serum levels are marked by *. Scale bar =1mm

### Effects of NT3 on afferent input reorganisation

Proprioceptive afferent terminals connect directly onto α-motor neurons through glutamatergic excitatory synapses. These connections are the primary source of vGlut1+ input directly onto motor neurons with contributions from Golgi tendon afferents and CST considered negligible in rodents ^72^. To quantify these connections the motor neurons innervating the ulnar aspect of the distal forelimb were retrogradely traced with Cholera Toxin beta subunit (CTb) and vGluT+ signal in close apposition to CTb+ motor neurons were quantified (Figure 16 A-C). The mature neuron marker NeuN was used to both accurately delineate the soma and to distinguish α-motor neurons from γ-motor neurons ^73^.

**Figure 16.**
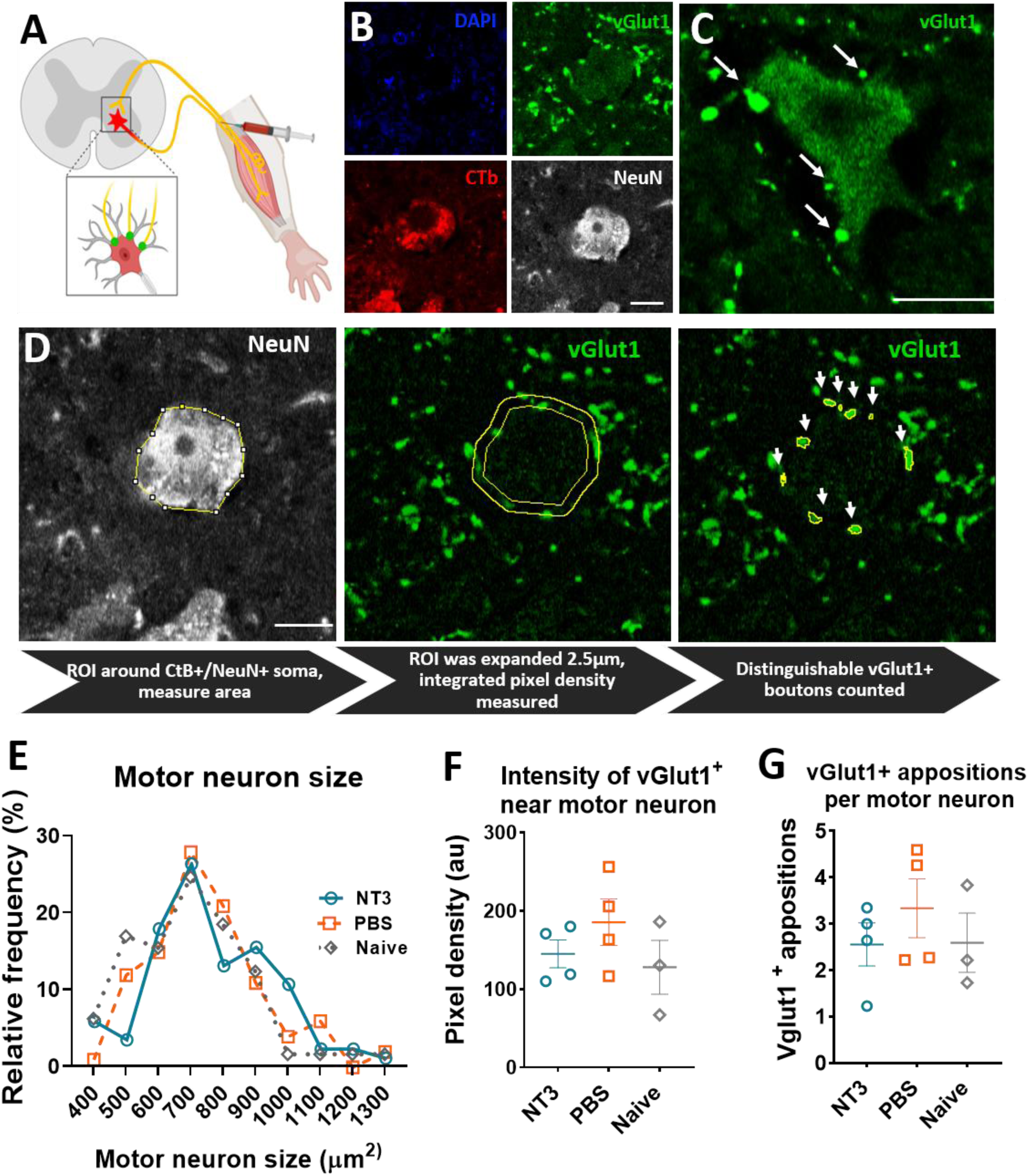
Assessing reorganisation of excitatory afferent input after injury and NT3 treatment. **A)** CTb was injected into the distal ulnar nerve, in order to trace motor neurons innervating distal forelimb muscles, and vGlut1 signal assessed as a proxy measure for afferent input onto this motor pool **B)** vGlut1 puncta was assessed on CTb+/ NeuN+ α-motor neurons within the ventral horn on the treated side. Individual vGlut1 puncta are mostly distinguishable from one another. **C)** Representative image showing the variation in size of vGlut1+ puncta. **D)** Overview of how analysis was performed. **E)** All CTb+/NeuN+ neurons within lamina IX of the C8-T1 segments which were analysed showed similar distribution of motor neuron soma size (Frequency histogram, with bin size of 100μm^2^). Average soma area remained unchanged following injury or NT3 treatment ( one way ANOVA F (2, 245) = 2.205, p=0.11, average size; NT3 =762±189μm2, PBS = 754±179μm2, naives = 702±183μm2) **F)** The integrated vGlut1+ pixel intensity in close proximity to the soma was tallied and normalised to the area of the band shown in D. There was no change in the vGlut1+ pixel intensity between the three groups (one-way ANOVA, F (2, 8) = 1.176, P=0.36). **G)** Individual vGlut1+ puncta in close proximity with CTb+/NeuN+ soma were counted. There was no difference between all three groups (one way ANOVA, F (2, 8) = 0.6063, P=0.56). Scale bar in B&D= 25μm, in C=5 μm. Animals per group: Naïve =3, NT3& PBS=4. Total number of motor neurons analysed: Naïve=65, NT3= 93 and PBS=108. Number of Puncta: Naïve=147, NT3= 239 and PBS=352

Successfully traced motor neurons followed a similar size distribution in all three groups (Figure 16D), and had an average soma size similar to previous reports of α-motor neurons^73^. Individual vGluT1+ boutons that were distinguishable from one another and in close proximity to the soma were counted. There was no change in the average number of vGluT1+ appositions between all three groups, suggesting no afferent sprouting occurred in these animals in response to injury.

Because the varying size of vGlut1+ immunoreactive puncta seen on the motor neuron perimeter suggested the possibility of two or more puncta not being distinguishable from one another, an additional method of quantification was performed . The raw integrated vGlut1+ pixel density was measured in a band 3μm thick surrounding the soma (Figure 16D), and was achieved by allocating an intensity to each pixel on a scale of 0-255, with 0=black/no signal and 255 = maximum brightness, and calculating the sum of all pixel intensities. This was normalised to the area of the region of interest and provides a complementary approach to quantifying the amount of vGlut1+ signal irrespective of number of puncta. There was no alteration in normalised raw integrated vGlut1+ pixel density following injury, shown in the PBS group compared to naives (Figure 16E). NT3 treatment also had no effect on integrated vGlut1+ pixel intensity when compared to either PBS or naïve. These results suggest that afferent input was not increased to these motor neurons following injury. Despite no significant changes between groups in either analysis, both methods revealed higher absolute puncta count or pixel density in the PBS treated animals compared to both NT3 and naïve group. This could be suggestive of injury induced afferent sprouting, however this analysis was only performed on a subset of animals prior to campus lockdown for COVID in 2020. Using the data collected for Figure 16G, if this magnitude of a difference did exist in real life it is calculated that a sample size of 18 per group would be required to detect a difference with p<0.05 and 80% power. This sample size exceeds the number of rats which remained in the study.

## Discussion

### Overview

In this study the potential therapeutic effects of NT3 on sensorimotor impairments in the upper limb after mid-cervical spinal cord injury were investigated. This built on previous work from our lab which showed forelimb function was improved following intramuscular treatment using AAV-NT3 after bilateral pyramidotomy ^57^. A mid cervical moderate contusion at the C5-C6 vertebral level generated sustained functional and electrophysiological impairments in the forelimb. Intramuscular AAV-NT3 treatment, initiated within the clinically feasible timeframe of 24h post injury, elevated NT3 protein levels in injected muscles and there is weak evidence to suggest this subtly reduced the deficit in locomotion. As anticipated from the previous pilot study, spinal cord injury (in the PBS group) induced hyperreflexia in H reflex of both forelimbs and this was sustained for 13 weeks. In addition, injury reduced Grip strength bilaterally and impaired the ability to grasp retrieve sugar pellets. Hyperreflexia was unaffected by NT3 treatment, in contrast to our previous study ^57^. NT3 was not able to improve grip strength or the ability to retrieve sugar pellets by 12 weeks. *Ex vivo* MRI revealed that the maximum extent of cavitation in the rostral direction was smaller in the rats that had received NT3. Histologically there was no strong evidence that sprouting of Ia afferent fibres occurred post injury or changes in afferent reorganisation as a result of NT3. Taken together the results suggests that the dose and delivery route of AAV-NT3 used was potentially beneficial to sensorimotor forelimb function but that these functional benefits were subtle and do not correlate with a reduction in hyperreflexia. They also suggest that NT3 as a monotherapy is unable to match its therapeutic effect following pyramidotomy ^57^, and as such is not able to overcome the additional damage to numerous motor pathways and grey matter populations caused by a contusion injury.

### Ex vivo imaging suggested that motor pathways other than the CST are spared after spinal cord injury

Contusive SCI lesions can be variable and reconstructing the lesion in therapeutic studies is essential to rule out the possibility of differing lesions between groups influencing outcomes. During processing for conventional histology, the delicate tissue around the lesion site can be affected from serial sectioning which can compress and deform the tissue. To avoid this, *ex vivo* MRI was used to generate T2 weighted scans with high resolution from which lesion size parameters like cavity volume can be automatically measured. Quantification of experimental SCI using MRI has been used previously and shown to adequately correlate with results generated using conventional histology ^74–76^. This study builds on previous analytical approaches involving manual quantification, by using an automated method to delineate the signals of interest such as hypointensities thought to correspond to cavitation and signal that is continuous with spared white matter at the extremities. This avoids variation that arises from selecting regions of interest manually and allows for a quicker quantification. Importantly, tissue can be processed later for routine immunohistochemistry, as performed in this study, without loss of antigenicity.

Ex vivo MRI of the lesion generated by a 225kdyn contusion revealed damage to several white matter tracts consistent with conventional histological findings ^77, 78^. Whilst individual white matter funiculi within the epicentre were not quantified, the dorsal white matter appears visually to be mostly disrupted whilst damage to the lateral and ventral white matter appears relatively spared . As a result it can be assumed that pathways associated with motor function located in these regions may have been partly spared.

The CST, conveying motor signals involved with fine motor movement, is split into three distinct portions descending in the dorsal, lateral and medial-ventral funiculi. A comprehensive study involving a <u>250 kDyn</u> injury to the same cervical region showed that the dorsal column projections are mostly damaged following a bilateral contusion but some ventral and dorsal-lateral CST fibres remain continuous through the lesion site ^79^. As *ex vivo* MRI showed a lack of putative spared white matter in the dorsal columns but some in the ventral and lateral white matter, loss of the dorsal CST with some sparing of ventral and dorsal lateral CST can be expected in this study.

The reticulospinal tract (ReST) arises in the reticular formation in the brain stem and descends the spinal cord in the ventral and ventrolateral funiculus in rodents ^80^. Given the regions of putative sparing, it can be assumed that some sparing of reticulospinal fibres has occurred. Propriospinal neurons are thought to be mostly intact after bilateral contusion injuries as they are uniformly distributed throughout the spinal cord lateral and ventral white matter ^81, 82^. The rubrospinal tract, which is essential for the arpeggio movement during reaching, was also partially spared as it descends in the dorsal lateral white matter ^83^ . Following injury the rubrospinal fibres reorganise by expanding their innervation of the forelimb motor pool, from predominantly distal muscles to include those of more proximally located muscles ^84^. These contribute to relay pathways allowing input from damaged supraspinal fibres, such as the CST and ReST, to transmit indirectly across the lesion and are contributing factors for recovery after SCI ^8, 85, 86^. The axons of short propriospinal neurons spanning the cervical region are susceptible to damage from contusion injury as they are located primarily on the white/grey matter boundary ^87^. This is most likely the case after 225kDyn contusion injury, where T2 weighted imaging reveals loss of the otherwise sharply demarcated boundary. In summary, *ex vivo* MRI indicated that various motor pathways other than the CST were spared by cervical contusion injury in the rat.

### Why was greater functional recovery not observed in this study?

Elevation of NT3 can induce neuroplasticity in several descending motor fibres. Functional recovery after thoracic contusion injury in mice occurs through reorganisation of propriospinal neurons promoted by NT3 ^48^. After treatment, propriospinal terminals were increased in the ventral horn, which partially restored descending neurotransmission to the hindlimbs. A potential mechanism suggested is NT3 mediated regrowth of motor neuron dendritic arbours, which would otherwise remain atrophic after injury, that enable a greater amount of propriospinal input to be received. Whilst that study focussed on hindlimb function it is plausible a similar mechanism would occur for forelimb function since short propriospinal neurons exist within the cervical spinal cord ^88, 89^. The lack of functional recovery in several forelimb tasks indicate that such relay pathways may not have formed in this study, or that retrograde transport to motor neurons was insufficient to cause an effect on dendritic arbours. The lesions generated at the T9 level in mice by Han, Ordaz *et al* (2019), have some sparing of the grey/white boundary region which is absent in this study. This would suggest damage to propriospinal axons was present and the subsequent disrupted transmission through the relay circuits accounting for the lack of functional recovery in cervical contusion compared to thoracic.

There are likely additional effects of NT3 which contribute to enhanced recovery. For instance, sensorimotor recovery with NT3 following pyramidotomy is unlikely to rely on propriospinal relay pathways spanning the lesion ^57^, since the injury generates complete ablation of the CST rostral to the level where propriospinal neurons connecting cervical spinal cord originate ^87^. Recovery was instead attributed to the normalisation of increased Ia afferent input onto motor neurons, possibly a result of NT3 mediated transcriptional changes within the DRG ^90^. Continual Ia afferent derived input into motor circuitry caudal to injury is essential for detour circuit formation and spontaneous limited recovery of locomotion ^6, 11^. Formation of these pathways may be enhanced through the remodelling effects of NT3 on afferent innervation onto forelimb motor neurons, primarily that of reducing inappropriate connections with heteronymous motor neurons ^57^. Afferent input onto motor neurons was thought to remain unchanged in this study but as it was not possible to determine the source of this input, i.e., from homonymous compared to heteronymous muscles, the result may include inappropriate connections. As a result, dysfunctional proprioceptive input may have been insufficient to stimulate these detour pathways to form.

It is also worth considering additional effects of NT3 associated with functional recovery which may not have been achieved in this study and help explain the lack of functional recovery. Sprouting of spared CST can be induced by elevated NT3 in the spinal cord or periphery. This could be achieved through diffusion across the blood-spinal cord barrier from circulating NT3 ^91^ or retrograde transport with motor neuron and secretion from the cell body, which can occur during development ^92^ Following this type of injury there is a limited number of CST fibres remaining caudal to the injury, and it is possible that these are incapable of generating enough collaterals to restore forelimb function. CST fibres rostral to injury will remain intact and sprout collaterals ^10, 44^, and these could contribute to recovery provided they form detour circuits involving descending propriospinal interneurons that control reaching ^8^. Given NT3 can enhance CST sprouting, it is plausible elevation would facilitate the formation of these relay connection. In the absence of any meaningful recovery there is a potential for; propriospinal neurons to be damaged in the contusion injury, disruption NT3 transport to the spinal cord parenchyma, or if transport did occur it was not abundant enough to influence CST plasticity.

### Why was Intramuscular NT3 unable to normalise hyperreflexia after contusion?

Following contusion injury animals developed hyperreflexia in proprioceptive circuitry caudal to the injury which persisted despite intramuscular NT3 treatment. Previously our lab has shown that intramuscular AAV-NT3 normalises the excitability of H reflexes following pyramidotomy from four weeks after treatment ^57^. Whilst it was not possible to verify these injections elevated NT3 in the ADM muscle, given its small size, it was assumed that it did based on the elevation found in another treated muscle, the Biceps Brachii. The ADM muscle, from which the H reflex was recorded, was injected with an equivalent dose totalling 3x10^9^ GC of AAV-NT3 in both studies. This was however achieved through different means; a single injection of 10μl in the pyramidotomy study compared to two 5μl injections in the present study. Another difference was the injection of a further 3x10^9^ GC of AAV-NT3 into the paw pad in the former study which was not performed here. It could be that diffusion of the viral vector throughout the paw contributed to higher titres within the ADM, or that treatment of these additional muscles, and their Ia afferents, contributes to the overall reduction of hyperreflexia.

A predominance for spasms in the forelimb flexors following pyramidotomy justified the decision in that study to treat only the flexors of the elbow and wrist. This is an important difference from the present study which, in the absence of any noticeable spasms during locomotion, treated both flexors and extensors as this was decided the best chance of restoring forelimb function. A consequence of this would be additional Ia afferents from forelimb extensors, such as *triceps Brachii,* being exposed and modulated by NT3. Whilst the H reflex incorporates a monosynaptic connection between Ia afferent and a homonymous motor neuron, it is in reality much more complex. Motor neurons also receive monosynaptic input from afferents originating in synergistic muscles termed heteronymous connections, for example between extrinsic wrist flexors and the motor neurons innervating muscles within the paw like the ADM ^93^. In addition afferents from antagonistic muscles form polysynaptic connections, via Ia afferent inhibitory interneurons, onto α-motor neurons ^94^. Hyperreflexia is likely to be caused by the sum of all input, from both homonymous and heteronymous muscles, converging on the motor neuron pool of the ADM muscle directly or via interneurons. Given that extensor muscles as well as flexors were treated with AAV-NT3 in this study, it is possible that the modulation of afferent input from extensors muscle counteracts any normalisation of hyperreflexia with treatment to flexor muscles.

There are several proposed mechanisms by which hyperreflexia arise such as; changes to chloride homeostasis, imbalance of excitatory and inhibitory signals converging on motor circuitry, alterations in dendritic arbours and innate increases in motor neuron excitability ^17, 25, 26, 57, 95, 96^. NT3 has been shown to modulate a number of these through a mix of *in vivo* and *in vitro* studies, which may not have occurred in this study and enabled persistent hyperreflexia.

NT3 can modulating dendritic branching and soma size of cortical and spinal neurons ^47, 97^. Such changes would alter electrophysiological properties of the cells to reduce the chance of depolarisation and reduce intrinsic excitability of the motor neuron. These properties have been detected in spinal motor neurons undergo following intramuscular AAV-NT3 ^98^. It is not possible to determine the excitability of the motor pool itself through H reflex testing ^19^. Motor threshold, an indication of motor axon excitability, can however reflect motor neuron excitability after SCI ^99, 100^. In this study there was no difference between groups in the intensity of stimulation needed to induce a motor axon response in the ADM muscle, although over time there was a gradual decrease in all three groups. This is likely caused by technical refinements made over time, such as less repositioning of the recording electrode which would reduce bleeding. Accordingly, we conclude there is no evidence for either SCI or NT3 treatment altering the excitability of motor axons, however additional electrophysiology would need to be performed to confirm this such as threshold tracking ^19, 101^ or measuring of persistent inward currents ^96^ .

Proprioceptive input onto motor neurons is glutamatergic and is essential for spontaneous motor recovery following SCI ^6, 72^. After bilateral CST lesions this input is increased, due to sprouting of the centrally projecting terminals of proprioceptive afferent fibres ^5, 57^. This appears contrary to what was found in this study after cervical contusion injury, where vGluT1+ signal in close apposition to motor neurons caudal to injury was unaltered. It is not clear if this reflects incompleteness of analysis or is a genuine finding. Afferent input can remain unaltered moderate cervical contusion ^102^, but does undergo transient reduction after thoracic staggered hemisections ^103^. Unfortunately, these studies used bulk analysis of vGlut1+ signal in distinct laminae or the entire grey matter respectively, rather than specifically quantifying input onto motor neurons. More work is required to determine whether contusion effects vGlut1+ signal on motor neurons. Whatever the case, is it possible additional excitatory inputs are altered which cause hyperreflexia to arise after injury?

It would be informative to assess the number of vGlut2+ and vGlut3+ boutons apposing motor neurons, which would provide a fuller understanding of how all excitatory inputs directly on motor neurons are altered after SCI and NT3 treatment. Direct excitatory drive to motor neurons have a widespread origin, arising from reticulospinal, vestibulospinal and rubrospinal tract as well as propriospinal and local interneurons ^104–106^. vGlut2 is present up to ten times more puncta per length of motor neuron membrane compared to vglut1 and is particularly abundant in the ventral horn after chronic staggered hemisection ^103, 107^. As of yet alterations in vGlut2 has not been explored in relation to hyperreflexia, however optogenetic activation of vGlut2+ interneurons can initiate tail spasms in a sacral preparation ^108^. Another possibility would be to extends analysis to include dendrites in addition to motor neurons. Proximal dendrites are the site of numerous vGlut1 synapses ^109^ and excitatory input into even distal portion of the dendritic arbour can generate excitatory post synaptic potentials in the soma^110^.

### Could central and peripherally acting effects of NT3 restore white matter?

An unexpected result from *ex vivo* MRI was the discovery of more putative white matter close to the epicentre in NT3 treated animals. This occurred despite no difference between the severity of injury given to each group. In order to automatically detect putative spared white matter the software takes the T2 weighted signal within a user-defined seed region, and then expands it to incorporate similar signal. As this seed region was one lateral funiculus 2.5mm rostral to the lesion an assumption was made that the white matter in this funiculus was spared. Whilst scar tissue surrounds the cavity and is likely represented by the hyperintense signal lining the cavity, it cannot be ruled out that this signal allocated as presumptive spared white matter may also incorporate scar or oedematous tissue or other pathology. A variety of possible explanations for NT3 mediated effects on white matter after injury are discussed below and include neuroprotection, axonal regeneration and remyelination. There is also chance this increase in putative spared white matter is a false positive, since these mechanisms might be expected to act equally throughout the lesion.

Whilst a neuroprotective effect of NT3 has been observed on axotomized motor neurons in neonatal rats ^111^, there was no increase in survival of adult corticospinal neurons following internal capsule lesion ^112^ or dorsal column injury ^37^. Cell loss of other neuronal populations, such as projection sensory fibres, may be prevented after NT3 is infused into the cavity after transection ^113^. Whilst from a technical perspective it would be interesting to see whether *ex vivo* T2W MRI is sensitive enough to detect a neuroprotective effect, such an effect is unlikely to occur in this study given the different injury mechanism and the 24h-delay between contusion and onset of treatment usingNT3.

One caveat to any neuroprotective role of AAV-NT3 delivered intramuscularly would be the delay from treatment at 24 hours post injury to raised NT3 in the lesion environment, which is likely to peak after apoptosis in the neuronal population peaks around 3 days ^114, 115^. Modulation of the immune response acutely after injury can halt the progressive secondary pathology resulting in reduced lesion sizes and improved functional outcome. There is modest evidence suggesting that NT3 can modulate the response of immune cells ^116^. Whilst serum NT3 levels were not elevated chronically, see below, there is the chance that a minimal increase in NT3 occurred after treatment and this is sufficient to affect the immune response. Blood-spinal cord barrier breakdown would be enhanced within the epicentre and access of NT3 to these zones may explain the spatially restricted increased in presumptive spared white mater in this region.

Another possibility is NT3 mediated signalling modulating axon regeneration and remyelination following injury. In addition to the effects of sprouting motor pathways previously mentioned, spinally applied NT3 is able to promote the growth of sensory axons after injury ^117, 118^, however regeneration following SCI is limited after treatment with a single growth factor and unlikely to be to such an extent that lesion cavity is impacted.

Regarding remyelination, oligodendrocytes express TrkC and exhibit both enhanced survival in culture and upregulation of myelin associated genes following NT3 application ^119–121^. A number of studies have shown beneficial effects of NT3 on remyelination following spinal cord injury^59^, such as increased myelination of regenerating neurons elongating into NT3 secreting fibroblast grafts ^122^. NT3 can also encourage migration of Schwann cells ^123^, which infiltrate from the periphery after SCI and contribute to remyelination ^124, 125^. Conversely, a recent study found no effect of NT3 as a monotherapy on myelination of regenerating axons after hemisection, however when combined with interleukin-10 was were modest effects ^59^. Recent studies have indicated that oligodendrocyte mediated remyelination plays no role in the recovery of hindlimb function following injury ^126^ although Schwann cell myelination may _do_ 127.

Slow release of NT3 from chitosan grafts appears to stimulate endogenous neurogenesis after a full transection which was associated with tissue bridges spanning the 5mm injury site, restoration of sensory and motor evoked potentials and functional hindlimb recovery ^128, 129^. Importantly, most aspects of this study have been independently repeated and the results replicated ^130^.

In summary, delayed treatment with AAV-NT3 may increase the amount of white matter near the lesion. Additional experiments would be required to identify the mechanism by which this occurs.

### Minimal elevation of NT3 in serum could indicate suboptimal expression of NT3 in muscle

Disappointingly, serum levels of NT3 were not elevated throughout all of the NT3 group. One rationale for NT3 driven recovery is through uptake by Ia afferents and/or motor neurons innervating muscle, which is maximised with high expression of NT3 within the muscle. Whilst elevation in the serum was not the primary target it is an unexpected result as previous studies using intramuscular delivery of NT3 result in serum levels consistently elevated at similar chronic time points ^57^. This discrepancy could reflect that maximal intramuscular expression of NT3 was not achieved in some animals, or the presence of other variables which limited its elevation above basal levels.

Our therapy is packaged in AAV serotype 1. This serotype exhibits a high tropism for muscle, estimated 10-100 times more than other serotypes ^131^ and is efficient at elevating serum levels following intramuscular delivery compared to AAV2-5 ^132^. Testament to this is its current use in a phase I/IIa trial for Charcot-Marie-Tooth Ia neuropathy where muscle transduction is essential (NCT03520751) ^133^. In the present study evidence of the high tropism for muscle, and potency of the CMV promoter used, is the 1500-fold increase in expression of NT3 in the *Biceps Brachii*.

As with other neurotrophins, NT3 is quickly eliminated from circulation with a half-life in plasma in the magnitude of minutes ^46, 134^. Following chronic infusion from subcutaneous osmotic pumps of NGF, another member of the neurotrophin family with a similar half-life in serum, the elimination half-life can be extended ^135^. Despite this, full elimination occurs within 24 hour and is suspected to be a similar time course for NT3. It is clear that for elevation to be sustained in serum long term, NT3 must be continually secreted and at a rate which surpasses the amount which is efficiently cleared.

For NT3 synthesised in muscle to be released into the circulation it must undergo extracellular secretion, diffusion through extracellular matrix surrounding myofibrils and finally pass through the endothelial wall. The endothelial monolayer is permeable to molecules with a molecular weight of up to 100kDa ^136^, much larger than NT3 at approximately 14kDa. No studies have explicitly looked at vascular permeability of NT3 but it is able to cross the tightly regulated blood brain barrier at a low rate ^134^,and presumably also able to at muscle capillaries. Capillary density varies massively depending of both muscle fibre type ^137^ and size ^138^ as a function of their oxidative capacity. Whilst fast type II fibres constitute the majority of the rat forelimb flexor and extensor muscles, there is variation within individual muscles associated with different capillary densities ^139, 140^. Since individual distal forelimb muscles were not always identified when injecting through the fascia, it is possible that certain muscles, or compartment of a muscle had high titre injected but had differing capacities to secrete NT3 into the circulation. In future, studies may benefit from either serum sampling at a middle time point to assess whether levels are ubiquitously elevated amongst the group, or analysis of all muscles treated to provide a total sum of NT3 being synthesised in the target tissue.

## Conclusion

This was the first study in which intramuscular AAV-NT3 was used in an attempt to treat forelimb sensorimotor deficits after bilateral spinal cord cervical contusion. Whilst the treatment was able to successfully recover some function of forelimb involvement in locomotion, it was subtle and did not alter the sustained hyperreflexia that occurred. This limited response to NT3 is likely a result of the highly complex injury with multiple pathological processes at play which is a major challenge when attempting to test therapies to ever more clinically relevant models. It is possible that AAV-NT3, a neuroplasticity-based therapy, may benefit being part of a combinational therapeutic intervention including intensive forelimb rehabilitation to maximise afferent input after injury and ensure functionally relevant connections are sustained.

## Acknowledgments

This research was principally funded by a Nathalie Rose Barr PhD studentship from the Spinal Research Trust (NRB117 to LDFM for JS-S). This project was also partly funded by the Medical Research Trust (S026053/1 to LDFM). We acknowledge the Penn Vector Core in the Gene Therapy Program of the University of Pennsylvania for production of the AAVs for this project. We thank Dr Diana Cash and Dr Eugene Kim (King’s College London preclinical imaging core) for their help with acquisition of magnetic resonance images. Thanks to Emily Burnside for training and advice regarding surgery. Thanks to Mohammed Ilyaas Javid for the pellet reaching boxes. Thanks to Zoe Hore for blind-coding the interventions.

## Contributions

JDS-S and LDFM designed the study. JDS-S and VM performed surgeries. ABS performed ELISA. LDFM obtained the funding. JDS-S and LDFM wrote the manuscript.

